# Efficient ultrasound-mediated drug delivery to orthotopic liver tumors – Direct comparison of doxorubicin-loaded nanobubbles and microbubbles

**DOI:** 10.1101/2023.09.01.555196

**Authors:** Pinunta Nittayacharn, Eric Abenojar, Michaela Cooley, Felipe Berg, Claire Counil, Amin Jafari Sojahrood, Muhammad Saad Khan, Celina Yang, Elizabeth Berndl, Marcin Golczak, Michael C. Kolios, Agata A. Exner

## Abstract

Liver metastasis is a major obstacle in treating aggressive cancers, and current therapeutic options often prove insufficient. To overcome these challenges, there has been growing interest in ultrasound-mediated drug delivery using lipid-shelled microbubbles (MBs) and nanobubbles (NBs) as promising strategies for enhancing drug delivery to tumors. Our previous work demonstrated the potential of Doxorubicin-loaded C_3_F_8_ NBs (hDox-NB, 280 ± 123 nm) in improving cancer treatment in vitro using low-frequency ultrasound. In this study, we investigated the pharmacokinetics and biodistribution of sonicated hDox-NBs in orthotopic rat liver tumors. We compared their delivery and therapeutic efficiency with size-isolated MBs (hDox-MB, 1104 ± 373 nm). Results showed a similar accumulation of hDox in tumors treated with hDox-MBs and unfocused therapeutic ultrasound (hDox-MB+TUS) and hDox-NB+TUS. However, significantly increased apoptotic cell death in the tumor and fewer off-target apoptotic cells in the normal liver were found upon the treatment with hDox-NB+TUS. The tumor-to-liver apoptotic ratio was elevated 9.4-fold following treatment with hDox-NB+TUS compared to hDox-MB+TUS, suggesting that the therapeutic efficacy and specificity are significantly increased when using hDox-NB+TUS. These findings highlight the potential of this approach as a viable treatment modality for liver tumors. By elucidating the behavior of drug-loaded bubbles *in vivo*, we aim to contribute to developing more effective liver cancer treatments that could ultimately improve patient outcomes and decrease off-target side effects.

## Introduction

Hepatocellular carcinoma (HCC) is the most common type of primary liver cancer and accounts for the majority of cases. It can be aggressive, have distinct characteristics, and pose its own challenges in terms of diagnosis and treatment(1). In addition to HCC, liver metastasis is a common and challenging complication of various malignancies, including colorectal, breast, and pancreatic cancer (2). Despite advancements in cancer therapies, current treatment options for both HCC and liver metastasis remain a clinical challenge due to factors such as limited drug delivery to the tumor site, systemic toxicity, and tumor heterogeneity (1,3). Conventional chemotherapy and passive drug delivery using nanomedicine for HCC and liver metastasis often face limitations, including inadequate drug penetration into tumor tissues, low drug accumulation in the tumor due to the tumor’s complex microenvironment, and off-target effects on surrounding healthy liver tissue (4). Accordingly, more effective treatment strategies are needed.

To overcome these challenges, there has been growing interest in ultrasound-mediated drug delivery using lipid-shelled microbubbles (MBs) as a promising strategy for enhancing drug delivery to solid tumors (5–9). MBs are small gas-filled colloidal particles with diameters between 1-10 µm (10). Since their U.S. Food and Drug Administration (FDA) approval as ultrasound contrast agents (UCAs), MBs have been widely used in off-label drug delivery and experimental pre-clinical applications (11,12). Generally, for drug-delivery approaches, ultrasound-mediated therapeutic delivery is achieved using targeted excitation of MBs with focused ultrasound at the region of interest. During insonation, MBs experience a process known as cavitation, which involves several mechanical and physical changes such as volumetric expansion, compression, oscillation, fragmentation, coalescence, dissolution, and/or abrupt collapse (6,13). These changes result in a temporary and reversible disruption of a nearby surface, allowing transient enhanced permeability across the cell membrane, vascular wall, and blood-brain barrier. This is known as sonoporation, and it allows for the released or co-administered therapeutic agents to extravasate into the target region.

In addition to co-administration of MBs and drugs, therapeutic agents can be loaded either in or on the shell of MBs. The MB shell is typically comprised of phospholipids, and well-known lipid conjugation strategies provide opportunities for functionalization and targeting of the MBs, making them outstanding theranostic agents (14–16). Several studies have demonstrated the potential of MBs for improving drug delivery to tumors and enhancing therapeutic efficacy (13,17–21). However, the clinical application of MBs in drug delivery is limited because MBs are typically too large to extravasate out of the circulatory system passively and seem to have difficulty penetrating the deep tissue layers. When MBs are destroyed by ultrasound (US) while within the vasculature, the therapeutic agents are released in the blood pool. There is a growing interest in using UCAs that show the same features as MBs in an acoustic field but are small enough to extravasate through leaky vasculature where there is increased permeability, such as with tumors and chronic inflammatory diseases (22). To improve tumor biodistribution and take advantage of the enhanced permeability provided by the leaky vasculature, the drug delivery complex should be less than 400 nm in size (14,23–25).

Interest in sub-micron bubbles, or nanobubbles (NBs), as vehicles for US-mediated drug delivery has significantly grown over the last decade due to the unique characteristics that make them well-suited for localized and targeted delivery of several therapeutic agents for various applications (26). NBs are generally considered to be UCAs with 200–600 nm diameters (27–31). They offer characteristics that could increase extravasation and diffusion through the extravascular space, including buoyancy, low density, high concentration, and high deformability (32). NBs have been shown to extravasate from the vessel and through the vascular endothelial barrier to reach tumor cells for targeted molecular imaging and drug delivery by the enhanced permeability and retention (EPR) effect (33–35). Our previous work has demonstrated the potential of NBs in improving drug delivery efficiency *in vitro* (30). We also showed that the drug loading capacity, acoustic performance, and therapeutic efficacy can be enhanced by simple deprotonation of doxorubicin (Dox.HCl) to hydrophobic doxorubicin (hDox) prior to loading it into lipid-stabilized NBs (31).

Both drug-loaded MBs and NBs have shown promising results *in vivo* (27,36–38). However, in the case of MBs, it is possible that at least some of the effects observed could be due to the presence of NBs in the generally heterogeneous sized bubble populations; prior studies have shown that commercial MBs formulations, such as Definity® have a substantial NB population (39–41). The delivery efficiency and short-term therapeutic efficacy have not been previously examined and compared directly between drug-loaded NBs and drug-loaded MBs *in vivo*. Understanding the differences in the kinetics of bubbles in tumors and the biodistribution of drug-loaded NBs and MBs is crucial for optimizing the drug delivery strategy and identifying the most effective bubble formulation for liver tumor therapy. Accordingly, this study investigated the effect of drug-loaded bubble size on therapeutic ultrasound (TUS) -mediated doxorubicin delivery to orthotopic liver tumors in an immunocompetent rat model. Bubble kinetics and biodistribution in the tumor and major organs were examined using NBs and size isolated MBs made from identical shell material and core gas. Acute damage induced by bubble cavitation and drug release was assessed in both tumors and the surrounding normal liver tissue 3 hours after treatment. By understanding the behaviors of drug-loaded bubbles *in vivo*, we aim to contribute to the development of more effective liver cancer therapies, which could eventually improve patient outcomes.

## Materials

Lipids including 1,2-dibehenoyl-sn-glycero-3-phosphocholine (DBPC), 1,2 dipalmitoyl-sn-Glycero-3-phosphate (DPPA), and 1,2-dipalmitoyl-sn-glycero-3-phosphoethanolamine (DPPE) were obtained from Avanti Polar Lipids (Pelham, AL), and 1,2-distearoyl-sn-glycero-3-phosphoethanolamine-N-[methoxy (polyethylene glycol)-2000] (ammonium salt) (mPEG-DSPE) was obtained from Laysan Lipids (Arab, AL). Octafluoropropane (C_3_F_8_) (Electronic Fluorocarbons, LLC, PA, USA). Doxorubicin hydrochloride (Dox.HCl) was from LC Laboratories (Boston, MA, USA) and was deprotonated using Triethylamine (TEA) to get deprotonated hydrophobic Dox (hDox). Daunorubicin (Dau) was purchased from Sigma Aldrich (Milwaukee, WI, USA). Propylene glycol (PG) was purchased from Sigma Aldrich (Milwaukee, WI). Glycerol was purchased from Acros Organics (Morris, NJ). Iscove’s Modified Dulbecco’s Medium (IMDM) with 10% fetal bovine serum and 1% penicillin-streptomycin, and trypsin-EDTA were purchased from Invitrogen (Grand Island, NY, USA). N1-S1 rat hepatoma cells were purchased from ATCC (Manassas, VA, USA). ApopTag® Peroxidase In Situ Apoptosis Detection Kit was purchased from Millipore (Billerica, MA, USA).

## Methods

### Preparation and purification of drug-loaded nanobubbles (hDox-NBs)

As previously described, NBs were loaded with deprotonated commercial Dox.HCl, which resulted in hDox (31). Dox.HCl was dissolved in a solution of chloroform and methanol (3:2, v/v), and it was then incubated overnight with TEA at a 1:3 molar ratio of Dox to TEA, deprotonating the sugar amino group as a result (42–45). After solvent evaporation, hDox powder was collected and stored in the freezer. Thin layer chromatography (TLC) was used to evaluate the status of hDox qualitatively. The materials were dissolved in tetrahydrofuran and spotted using a microcapillary on a silica gel TLC plate (TLC silica gel 60 F254, Merk, Darmstadt, Germany). The plates were exposed to UV light after being developed in a mobile phase made of dichloromethane, methanol, formic acid, and deionized water (82:24:2:1, v/v).

Preparation of drug-loaded NBs involved encapsulating hDox in lipid-shell stabilized octafluoropropane (C_3_F_8_) bubbles, which was previously described (31). In brief, a combination of lipids, including DBPC, DPPA, DPPE, mPEG-DSPE, and 0.2 wt. % hDox, was dissolved in PG. The lipid solution was then mixed with glycerol and phosphate buffer saline (PBS), and the air inside a 3-mL vial was replaced with C_3_F_8_. The vial was subsequently shaken for 45 seconds on a VialMix shaker (Bristol-Myers Squibb Medical Imaging, Inc., N. Billerica, MA) to promote bubble self-assembly. The NBs were extracted from the mixture by inverting the vial and subjecting it to centrifugation at 50 g for 5 minutes. The drug-loaded NBs were separated from free drugs by passing the mixture solution of drug-NBs and free drug through a Sephadex G-25 in the PD-10 desalting column. The NBs were eluted through the column with PBS/PG/glycerol (10.3:15.6, %w/v) solution and the first 3 mL fraction was collected for further experiments.

### Isolation of hDox-loaded microbubbles (hDox-MBs)

MBs with a size ranging between 0.8-1.5 µm were obtained from the activated bubble vial, which contained a mixture of MBs and NBs, through an adapted differential centrifugation method (45,46). The bubble mixture from two vials was transferred to a 50 mL centrifuge tube and diluted with PBS containing 10% (v/v) of glycerol and PG to obtain a total volume of 30 mL. The solution was then loaded into a 30-mL syringe and centrifuged at 300 g for 10 minutes. The liquid suspension, or infranatant, was discarded, and the MBs that formed a cake on the syringe plunger were re-dispersed in 30 mL of PBS. The re-dispersed MBs were then centrifuged at 70 g for 1 minute to eliminate large bubbles. The infranatant was then centrifuged at 300 g for 10 minutes to obtain the cake containing MBs with the desired size. The MB cake was re-dispersed in 2 mL of PBS, and the previous purification methods were employed for equivalent experiments to remove free drugs from drug-loaded MBs.

### Determination of drug-loading content

The drug-loaded NB and MB solutions were subjected to ultrafiltration using a Vivaspin® 20 unit (Sartorius, Gottingen, Germany) with a molecular weight cut-off of 50,000 Da to remove any free drug (31). The process involved centrifuging the solution at 4,000 rpm for 50 minutes. The resulting hDox-NBs solution was then lyophilized, weighed, and dissolved in a mixture of PBS and methanol (1:1, v/v). The fluorescence of hDox was measured using a TECAN plate reader (Infinite M200, San Jose, CA, USA) with an excitation of 495 nm and an emission of 595 nm. The amount of encapsulated drug in both NBs and MBs was determined by a calibration curve generated using known amounts of the drug dissolved in PBS and methanol (1:1, v/v). The same methods were used to perform equivalent experiments with hDox-MBs. NB and MB doses were normalized by total Dox loading dose for the following *in vivo* studies. Purified drug-loaded NBs and MBs were diluted with PBS to achieve a final administration Dox dose of 375 mg/kg in rats.

### Characterization of size and concentrations

The size distribution and concentration of buoyant hDox-NBs and hDox-MBs were measured using resonant mass measurement (RMM) (Archimedes, Malvern Pananalytical Inc., Westborough, MA, USA) with a calibrated nanosensor (100 nm – 2,000 nm) in three replicates. The sensors were pre-calibrated using NIST traceable 565 nm polystyrene bead standards (ThermoFisher 4010S, Waltham MA, USA). The hDox-NBs/MBs were diluted 1:100 with PBS, and 1,000 particles were measured for each trial. In addition to the RMM technique, Dynamic Light Scattering (DLS) analysis of hDox-NBs and hDox-MBs was also performed using a nanoparticle size analyzer (90Plus, Brookhaven, NY, USA) for comparison. The hDox-NBs/MBs were diluted 1:1,000 with PBS. For each sample suspension, five DLS measurements were performed with a fixed run time of the 20 sec. The scattering angle was set at 90°.

### Orthotopic hepatocellular carcinoma (HCC) rat model

The N1-S1 (48–50) rat hepatoma cell line was cultured in Iscove’s Modified Dulbecco’s Medium (IMDM), supplemented with 10% fetal bovine serum and a 1% penicillin/streptomycin mixture. Prior to tumor inoculation procedures, the viability of the cells was tested using the MoxiCyte Viability Kit (ORFLO, Ketchum, ID, USA), which confirmed cell viability of over 95%.

Male Sprague Dawley (SD) rats weighing 400 g and aged 3 – 4 weeks were procured from Case Western Reserve University’s animal research center. These rats were handled according to the Institutional Animal Care and Use Committee (IACUC) guidelines at Case Western Reserve University and given a week to acclimatize before tumor inoculation. The N1-S1 rat hepatoma model was initiated by injecting 12 x 10^6^ N1-S1 cells suspended in Matrigel into the liver of the rats via mini-laparotomy under general anesthesia. Before the procedure, the upper abdomen was shaved and cleaned using betadine and 70% ethanol solutions. A 2-cm midline laparotomy was conducted, and the left lobe of the liver was exposed. The N1-S1 cells were introduced into the subcapsular area, after which the muscle layer was sealed with absorbable sutures, and the skin layer was sealed with non-absorbable monofilament sutures. To prevent incision site infections, the animals were carefully monitored every day. Tumor growth was evaluated by an Acuson S3000 (Siemens Healthineers. Erlangen, Germany) with a 18L6 linear array probe (frequency 6 – 18 MHz) using B-mode and color doppler. Weekly scans were performed to detect and quantify tumor progression. All tumors were evaluated both in sagittal and coronal planes and volumes were estimated based on the standard clinical ellipsoidal equation(51).

### Perfusion dynamic study

Contrast-enhanced ultrasound scans were performed to quantify the kinetics of hDox-NBs and hDox-MBs in the tumors using the same scanner and transducer as above. Rats were anesthetized and placed in the face-up position, and the ultrasound probe was placed longitudinally to the axis of the animal body to visualize the liver. A comparison of contrast-enhanced ultrasound images in liver tumors was performed by administering 1 mL of either hDox-NBs or hDox-MBs via the tail vein and acquiring images using the Cadence contrast pulse sequencing (CPS) protocol (52–54). Bubble doses were normalized by total Dox loading per injection at 327.75 µg/kg. Raw data format images were acquired for 5 sec before the injections, and nonlinear CPS mode was used to image the change in tissue contrast density after the NBs or MBs injections. The rats were imaged continuously for 20 min at 8 MHz with a 18L6 probe, 0.1 MIF, and a frame rate of 1 frames/s. MIF refers to the maximum of the mechanical index measured at the active focal zone (55).

### Experimental protocols and hDox-NBs and hDox-MBs delivery

After tumor implantation, rats were randomly divided into five groups (n = 5 per group), and tumors were allowed to grow until they reached a size of at least 80 mm^3^. Once the tumors were of appropriate size, the animals (n = 5) received treatment as follows: 1) hDox-NBs only; 2) hDox-NBs with the combination of low-frequency ultrasound(hDox-NB+TUS); 3) hDox-MBs only; 4) hDox-MBs with the combination of low-frequency therapeutic ultrasound (hDox-MB+TUS). Animals (n = 2) without any treatment were used as a control group. The rats were anesthetized with 1 – 2% isoflurane and 0.5 – 1 L/min oxygen and were given a dose of hDox-NBs or MBs (Dox dose: 327.75 µg/kg) via the tail vein. For the group with TUS, 2 min after bubble administration, the tumors were exposed to TUS with a 1cm^2^ effective radiating area transducer (Sonicator 740, Mettler, Ca, USA) at 3 MHz, 2.2W/cm^2^, 10% duty cycle for 5 minutes. A peak negative pressure (PNP) amplitude of 0.25 MPa was estimated at these parameters. Three hours later, the rats were euthanized, and organs, including the liver, lungs, kidneys, heart, and spleen, were removed, washed with saline, and blotted. These organs were used for the following *ex-vivo* experiments. The experimental scheme is shown in Figure 1.

**Figure 1.**
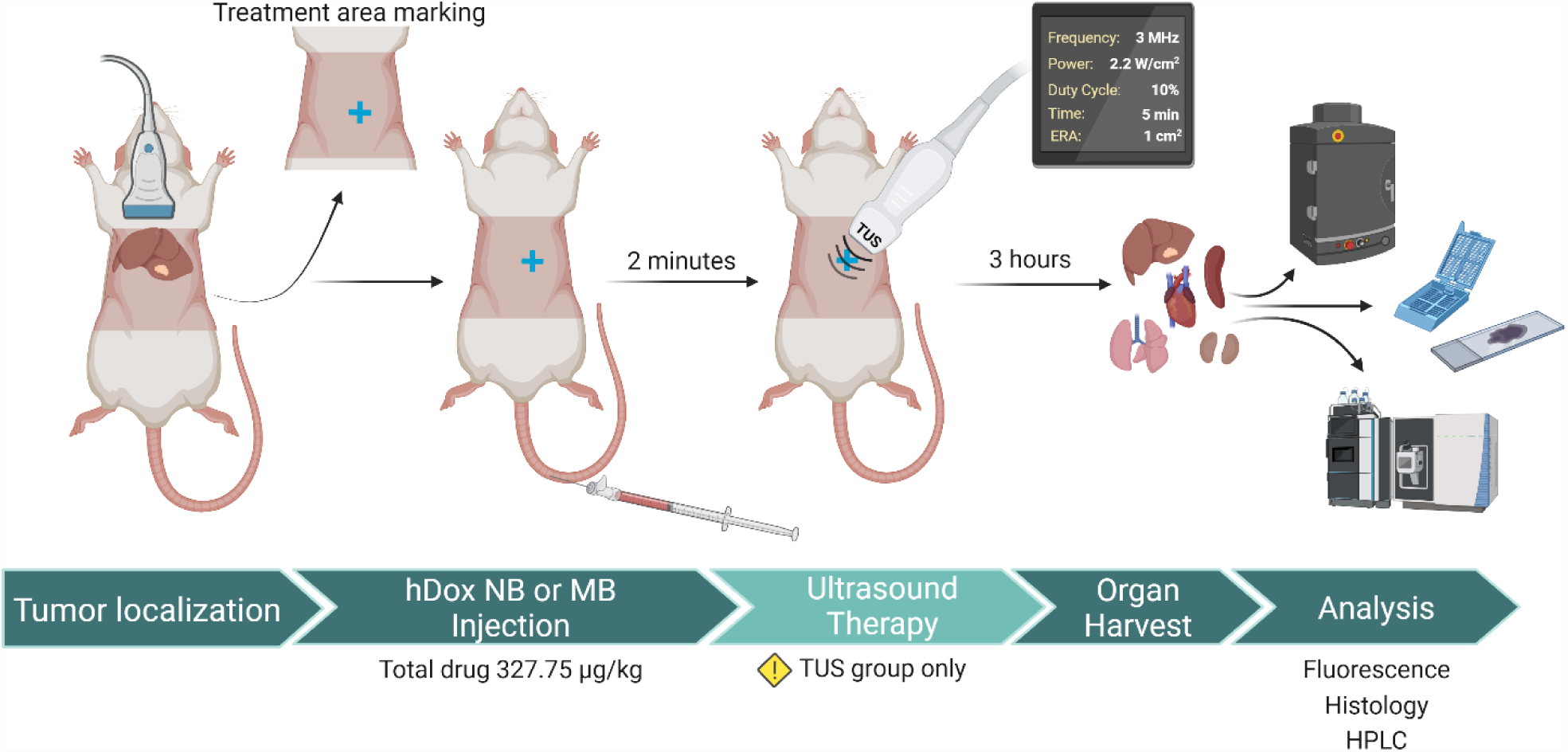
Schematic diagram of the *in vivo* experimental approach. Prior to bubble administration, tumors were localized and marked using B-Mode US. hDox-NBs or hDox-MBs was intravenously injected via tail vein and all animals (n = 5) received the same total drug administration (Dox dose at 327.75 µg/ kg). For the group with TUS, 2 min after injection, tumors were irradiated with TUS at 3 MHz, 2.2 W/cm^2^,10% DC for 5 min. 3 hours later, animals were euthanized, the tumor and organs were harvested for analyses, including 1) optical fluoresce imaging; 2) immunohistochemistry; 3) tissue extraction, and LC/MS.

### Biodistribution study

The hDox fluorescence intensity in each organ was measured using an optical fluorescence imaging system (CRI Maestro, Caliper Life Science) with a blue excitation/green emission filter (445-490 nm/long pass 580 nm) (56,57) and an exposure time of 500 msec. Organs from rats without Dox were used as a baseline subtraction. The fluorescence images were saved in JPEG format, and ImageJ software (freeware available from NIH) was used to calculate the mean fluorescence intensity of each organ.

### Histopathological analysis

After euthanasia, the liver, kidneys, spleen, heart, and lungs were harvested. Longitudinal sections at the periphery of the tumors were preserved in a 4% paraformaldehyde solution for 24 hours, then transferred to 30% sucrose and kept in this solution for 48 hours (56,57). The tissues were embedded in an optimal cutting temperature solution and sectioned into 7 μm slices using a cryostat. The slices were stained with terminal deoxynucleotidyl transferase dUTP nick end labeling (TUNEL) and scanned using a digital slide scanner (Nanozoomer S60, Hamamatsu Photonics K.K., Japan). The positive pixel count algorithm in ImageScope software was used to analyze the images (58,59). In this algorithm, each pixel within a defined region of interest was labeled as weak, moderate, or strong, and a markup image was generated. The percentage of the apoptotic area was calculated by dividing the total number of combinations of moderately and strongly labeled pixels by the total number of pixels, including the negative pixels.

### Dox extraction from tissue

To quantify the amount of Dox accumulation in the tumor, the tissues were extracted following the protocol in the previous work (30). The tumors were washed in PBS and then homogenized in water at a ratio of 10 mL per gram. The resulting tissue homogenate was mixed with acidified isopropanol (0.75 N HCl) at a ratio of 1:4 (v/v), and the mixture was thoroughly combined and incubated for 3 hours at a temperature of 20°C. Following this, the samples were vortexed and centrifuged at 21,130 g for 20 minutes, and the supernatant was collected.

### LC/MS-based detection and quantification of doxorubicin in tissue samples

Detection of Dox was achieved using a Linear Trap Quadrupole (LTQ) linear ion trap mass spectrometer (MS) (Thermo Scientific) equipped with an electrospray ionization (ESI) interface (Thermo Scientific) and coupled to an Agilent 1100 HPLC (Agilent Technologies). Tissue extracts were injected into a reverse-phase C18 X-Bridge HPLC column (2.1 × 100 mm, 3.5 μm) (Waters). Chromatographic separation of the drug was achieved with a linear gradient of acetonitrile in water containing 10 mM ammonium formate and 0.1% formic acid (v/v). The gradient was developed from 10% to 100% over 10 min at the isocratic flow rate of 0.3 mL/min. The HPLC eluent was directed into the MS via the Electrospray Ionization (ESI) probe operated in the positive ionization mode. Parameters of ionization and detection were tuned with the synthetic standard of Dox to attain the highest possible sensitivity. For quantification, the analyzed samples were supplemented with a known amount of daunorubicin, which served as an internal standard (IS). Dox and Dau were detected by selected reaction monitoring (SRM) mode with the following ion transitions 544.2 → 397.4 and 528.2 → 363.3, respectively. The amounts of the drug in the analyzed samples were calculated according to a calibration curve, which correlated the ratio of the areas under SRM ion intensity peaks and molar ratios of the drug and the IS.

### Statistical analysis

Unless explicitly stated otherwise, the data are reported as the mean value ± the standard deviation. The statistical analysis of the data was performed using GraphPad Prism9 software, with ANOVA and Tukey multiple comparison tests. In general, a p-value less than 0.05 was considered to be statistically significant unless otherwise specified.

## Results

### *In-vitro* hDox-NBs and hDox-MBs characterization

Dox.HCl was deprotonated, resulting in the formation of the deprotonated hydrophobic Dox (hDox). The state of hDox was verified using a TLC plate, as shown in Figure S1. hDox-loaded NBs and MBs were successfully prepared and purified for removing free drug using our standard preparation process (31). The average diameter and concentration of hDox-NBs and hDox-MBs were determined using RMM. A representative histogram plot showing the concentration distribution of buoyant particles (bubbles) of hDox-NBs and hDox-MBs is shown in Figure 2A. The hDox-NBs had an average diameter of 280 ± 123 nm, while the hDox-MBs had an average diameter of 1104 ± 373 nm. The average concentration of hDox-NBs and hDox-MBs was 4.06x10^10^ ± 6.9x10^8^ and 3.18x10^9^ ± 2.9x10^8^ particles/mL, respectively. Most of the samples in a purified solution consisted of buoyant particles, accounting for 96.5% of MBs and 98.5% of NBs. Non-buoyant particles, likely a combination of micelles and lipid aggregates, were also present in both formulations but mostly removed during the purification process. Additionally, the size of both NBs and MBs was measured using DLS, the current gold standard for sizing nanoparticles. DLS results were consistent with RMM, where the average hDox-NB diameter was 273 ± 39 and 417 ± 63 nm, as a number and intensity-weighted while the hDox-MB diameter was 935 ± 274 and 1524 ± 211 nm as number and intensity-weighted averages (Figure 2B). The drug loading content (DLC) of hDox in particles, including both bubbles and non-buoyant particles, of hDox-NBs and hDox-MBs after purification was determined by centrifuge filtration and presented as DLC in µg/mL. The DLC in hDox-NBs and hDox-MBs was found to be 151.97 ± 7.6 µg/mL and 131.11 ± 15.3 µg/mL, respectively (Figure 2C). For comparison, the total hDox dose in the following *in vivo* experiments was normalized at 327.75 µg/kg. Bubble properties after hDox dose normalization are shown in Figure 2D. hDox-NBs were also characterized in a tissue-mimicking agarose phantom (60) using US parameters similar to the perfusion dynamic study to ensure their acoustic property prior to administration in mice (Figure S2).

**Figure 2.**
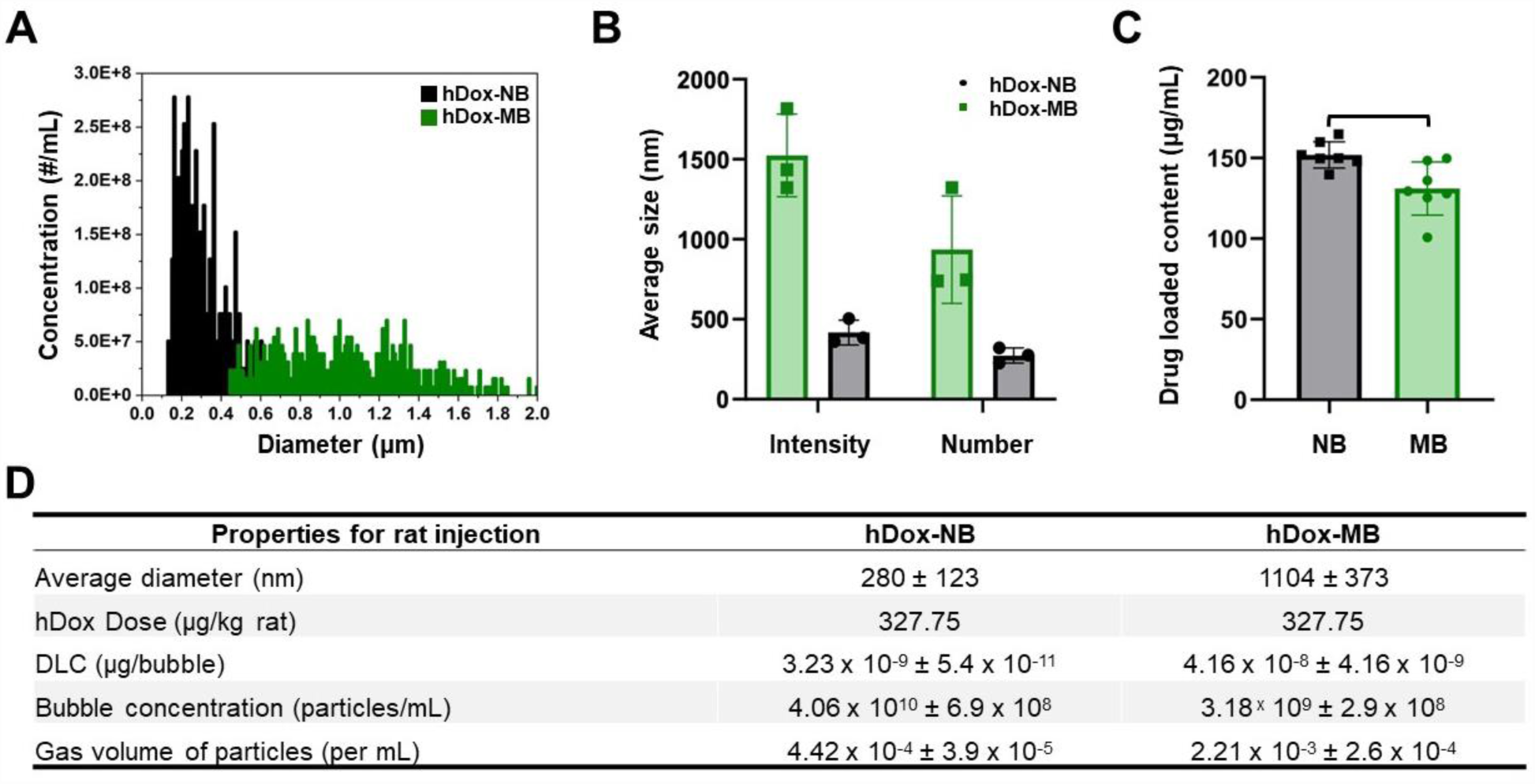
hDox-NBs and hDox-MBs were characterized using a resonant mass measurement (RMM) instrument, giving representative histogram of size and concentration for the sample’s buoyant particles (bubbles) (A); DLS measurements were performed to determine the diameter of the particles. Both RMM and DLS results were comparable (B); hDox-loading concentration per mL of bubble solution (C); Summary of bubble properties after normalization based on hDox dose for rat injection (D). Error bars represent the standard deviations.

### Perfusion dynamics of hDox-NBs and hDox-MBs in a rat tumor

To achieve the maximum cavitation efficiency from the bubble, attenuation measurements at several injection volumes were carried out prior to the application of treatment. hDox-MBs and hDox-NBs were imaged using nonlinear CPS mode at 8 MHz with an 18 MHz probe. Tissue attenuation was observed in hDox-MBs at a 1.5 mL injection volume as visualized by the decrease in signal at the beginning of the acquisition and then increasing over time, as shown in the time-intensity curves after hDox-MBs administrations shown in Figure S3A. On the other hand, hDox-NBs showed no attenuation at an injection volume of up to 2 mL, as depicted in Figure S3B. The dynamics of hDox-NBs and hDox-MBs within the tumors were then evaluated. Rapid signal enhancement was observed in tumors imaged with either hDox-NBs or hDox-MBs, reaching peak intensity within 0.8 to 1 min post-injection. The purified bubbles exhibited higher peak intensity and longer circulation time for hDox-MBs (Figure 3A). The representative contrast-enhanced ultrasound images of bubbles in the tumor at different time points are shown in Figure 3B. The summary of quantitative kinetic parameters of the tumor obtained from the time-intensity curve (TIC) is shown in Table S1. The peak enhancement of hDox-MBs was 12.5% higher than hDox-NBs. The half-life of hDox-MBs was 7 min, while that of hDox-NBs was 5 min. Based on the tumor perfusion dynamics of hDox-NBs and MBs, a potential starting time point for TUS irradiation and treatment duration was determined. Tumors were exposed to TUS at 2 min post-injection, and the irradiation continued for 5 min. By choosing this timepoint, TUS exposure started when hDox-MBs showed 82.81% of the peak enhancement and the hDox-NBs showed 79.32%. This would allow sufficient time for the NBs to extravasate via passive diffusion (36,37,61).

**Figure 3.**
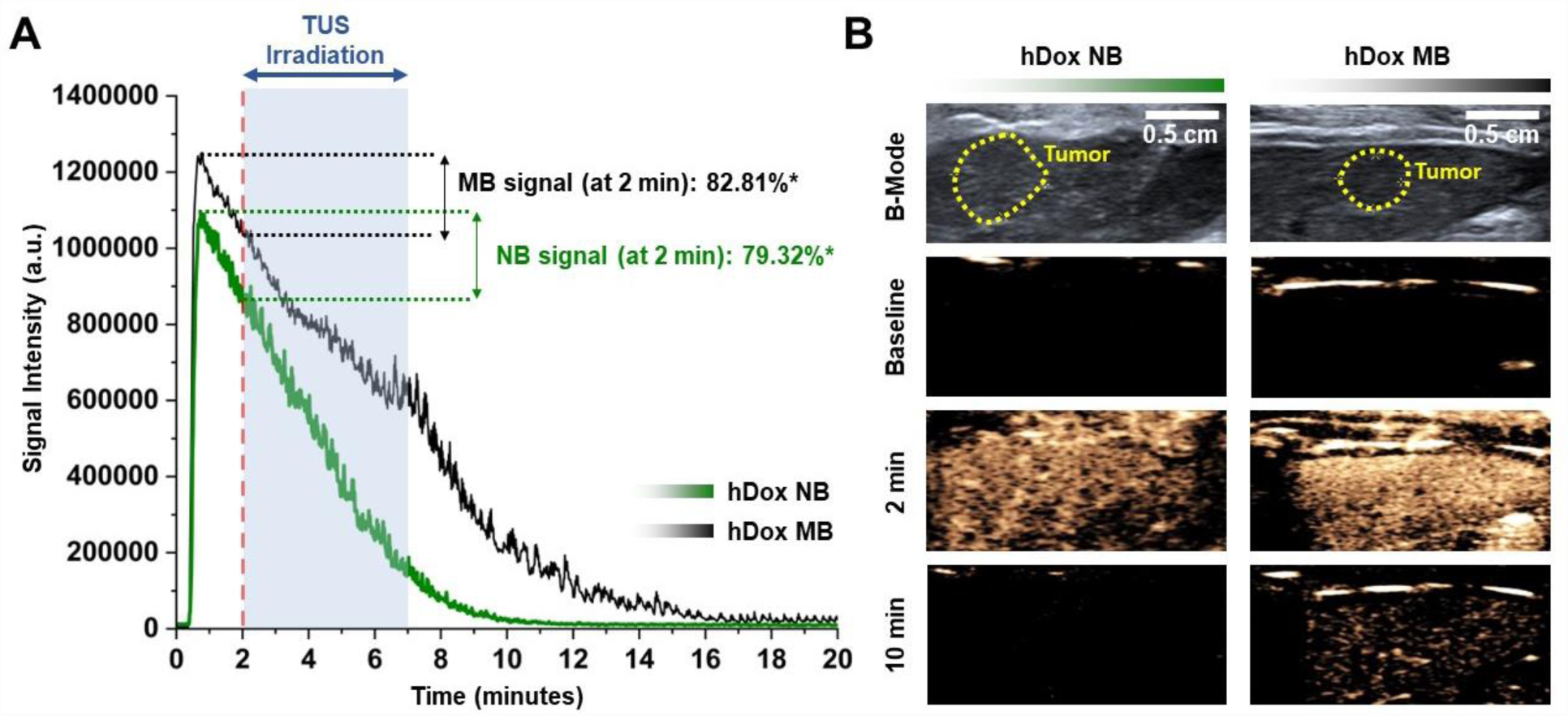
Time-intensity curves of the tumor after bubble administration. *Represents the percentage of NBs/MBs over the peak enhancement (A); Representative ultrasound images of tumor (yellow dashed line) in the liver. Prior to treatment, the location of the tumor was imaged using clinical ultrasound in B-mode. NB/MB enhancement of the tumor in the liver was imaged at 8 MHz with an 18 MHz probe, 0.1 MI (B).

### Biodistribution analysis

The biodistribution of Dox was assessed in Sprague Dawley rats (n = 5) following intravenous injection of hDox-NBs and hDox-MBs at a consistent Dox dose of 327.75 µg/kg. Three hours after treatment, rats were euthanized, and the organs were collected for *ex vivo* fluorescence imaging of Dox distribution using an optical fluorescence instrument. Representative images of Dox fluorescence in livers and tumors are shown in Figure 4A. The mean fluorescence intensity of Dox in tumors and organs was quantified by ImageJ software (Figure 4B). The highest Dox signal in each treatment group was found to be higher in tumors and the liver compared to other organs. However, the difference in Dox signal of each organ among the treatment group was not statistically significantly different. The Dox signal in the heart and lungs was negligible, while most signals were found in the spleen. Corresponding representative fluorescence images of organs are shown in Figure S4. In general, the hDox-NB group exhibited a higher Dox signal within the tumor when compared to hDox-MB group, and this signal was further amplified with the application of TUS. The treatment with hDox-NB+TUS led to a notable 26.7% increase in Dox signal within tumors in comparison to the hDox-MB+TUS treatment. Interestingly, when combined with sonication, hDox-MB resulted in a 2.2-fold increase in drug levels within the normal liver when compared to hDox-MB alone. In contrast, the utilization of hDox-NB in conjunction with TUS, resulted in a 2-fold *decrease* in Dox levels in the normal liver compared to hDox-NB without TUS. Notably, both hDox-NB+TUS and hDox-MB+TUS led to a reduction in Dox signal by 1.8 and 1.5-fold, respectively, within the spleen. Regarding the kidney, the application of ultrasound to the tumor region following hDox-NB injection yielded a 1.8-fold increase in Dox signal, whereas the increase for sonicated hDox-MBs was comparatively lower at 1.4-fold compared to bubbles without sonication.

**Figure 4.**
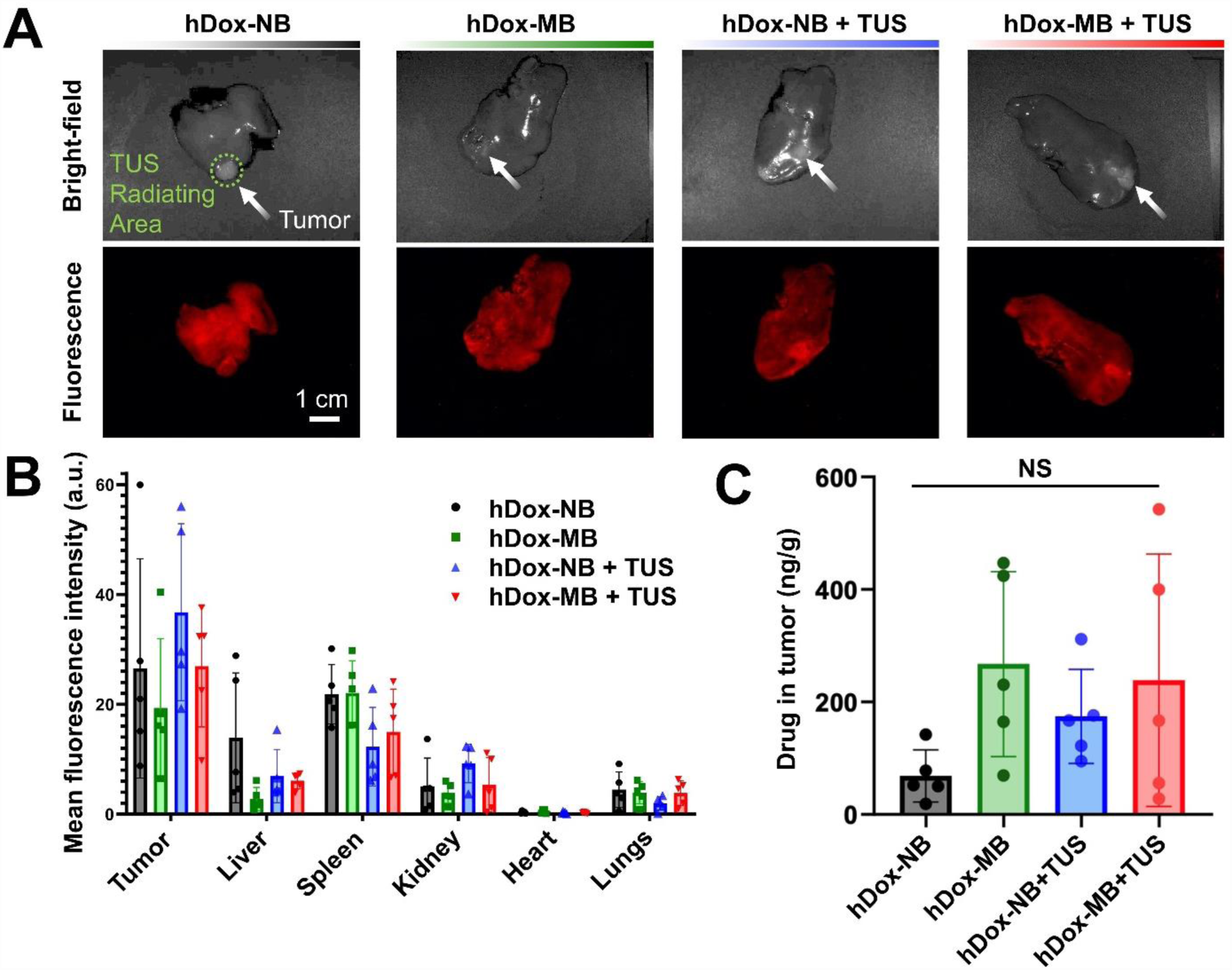
Representative optical fluorescence images of the liver and tumor from each treatment group showing hDox distribution at 3h after administration of hDox-NBs and hDox-MBs at consistent Dox dose. Bright-field images and fluorescence are shown in the top and bottom rows. The green-dash line represents the TUS effective radiating area. The tumor location on the liver is indicated by the white arrow (A); Average fluorescent intensity of hDox in rat tumors and organs including liver, spleen, kidney, heart, and lung. Organs without hDox were used as baseline signals (B); Dox accumulation in the tumor tissue at 3h time points after administration of hDox-NBs and hDox-MBs with consistent Dox dose. Dox levels in tumors were determined by acidified isopropanol extraction from tumor homogenates. Aliquots from five rats per group were analyzed by LC/MS (C).

Tissue extraction was performed to quantitatively assess the actual accumulation of Dox in tumor tissue. Tumors were excised from rats and Dox were extracted from the tissue using the previously described extraction protocol (30). Dox accumulation was quantified by LC/MS analysis. The amount of hDox in tissue was calculated by a calibration curve with known amounts of Dox and Dau dissolved in the same solvent solution used for tissue extraction (Figure S5). As shown in Figure 4C, rats treated with hDox-NB (68.32 ± 41.5 ng/g) and hDox-NB+TUS (174.29 ± 74.7ng/g) show a slightly lower accumulation of Dox in tumors compared to the group of hDox-MB (267.36 ± 147.0 ng/g) and hDox-MB+TUS (238.5 ± 200.6 ng/g). The difference is not statistically significant. However, most notably, the application of ultrasound to tumors resulted in a near three-fold increase in tumor Dox levels when hDox-NBs were used. In contrast, the application of TUS to hDox-MBs resulted in a decrease in overall tumor drug levels.

### Apoptotic analysis

To further investigate the therapeutic efficiency by assessing the acute damage in the tumor and surrounding liver tissue, immunohistochemistry (IHC) was performed on tumor sections. Representative TUNEL slides from tumors in the liver are shown in Figure 5. The brown-colored stain area indicates the apoptotic cells with DNA damage while nuclei in normal tissue were counterstained with hematoxylin and presented in a deep blue to purple color. The tumor region is delineated by the red dashed line. 5x and 40x images from the liver and tumor are also shown, respectively. In the absence of TUS, the hDox-NB group (Figure 5A) showed a higher apoptotic area in the tumor compared to hDox-MBs (Figure 5B). Strikingly, despite similar total drug accumulation, a significantly increased area of apoptotic cell death in tumor and fewer off-target apoptotic cells in the normal liver were found in the treatment with hDox-NB+TUS (Figure 5C).

**Figure 5.**
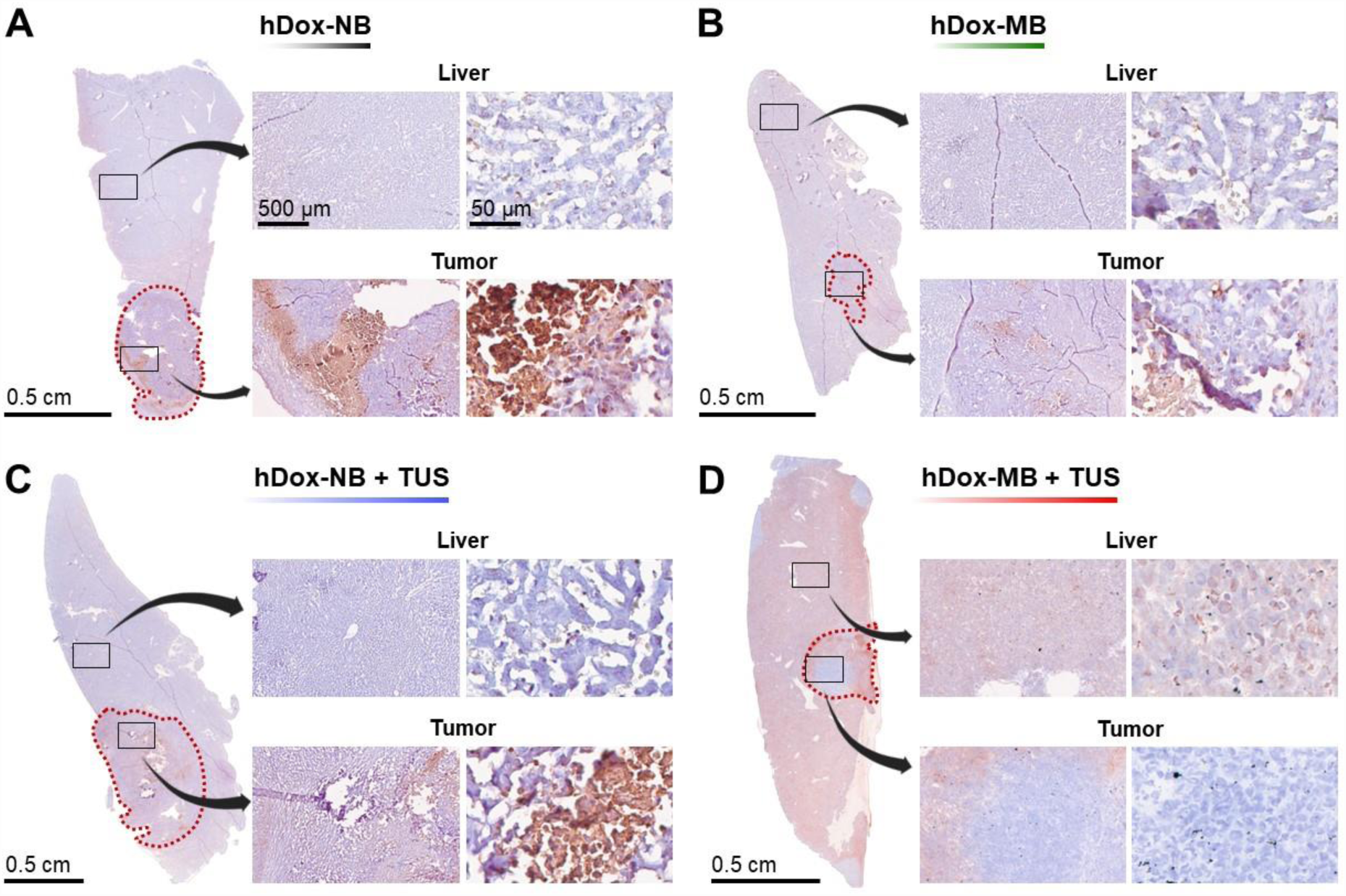
Representative images of TUNEL staining of the liver with tumor tissues from rats treated with hDox-NB (A); hDox-MB (B), hDox-NB+TUS (C); hDox-MB+TUS (D). Brown colored regions are indicative of apoptotic cells with DNA damage. The red dot line indicates the tumor boundary. 5x and 40x magnifications of tumor and liver areas are shown, respectively. The observed cell death in the liver for the hDox-MB+TUS group was widespread, while the hDox-NB+TUS group showed a more localized effect within the tumor.

In contrast, the cell death area was clearly seen everywhere throughout the liver from the group of hDox-MB+TUS and only a few were found in the tumor (Figure 5D). Using the Positive Pixel Count algorithm in ImageScope software, apoptotic cell numbers were quantified within these slides, showing the percentage of apoptotic cells in the region of interest (Figure 6A). For hDox-NBs, the apoptotic area in the tumor and liver was 9.10 ± 3.6% and 2.96 ± 0.4%, respectively. In contrast, hDox-MBs exhibited 2.48 ± 0.7% and 2.94 ± 0.2% apoptotic area in the tumor and liver, respectively. When TUS was applied, hDox-NB+TUS displayed a higher apoptotic area in the tumor compared to hDox-MB+TUS (17.01 ± 10.6% vs 5.92 ± 2.4%) and fewer off-target apoptotic cells in the normal liver (3.32 ± 1.7% vs 8.81 ± 2.6%). Furthermore, the tumor-to-liver apoptotic ratio was elevated 9.4-fold following treatment with hDox-NB+TUS compared to hDox-MB+TUS (Figure 6B). This is consistent with the LC/MS data shown above, where tumor drug levels were reduced following MB sonication, but were significantly increased following NB sonication in tumors.

**Figure 6.**
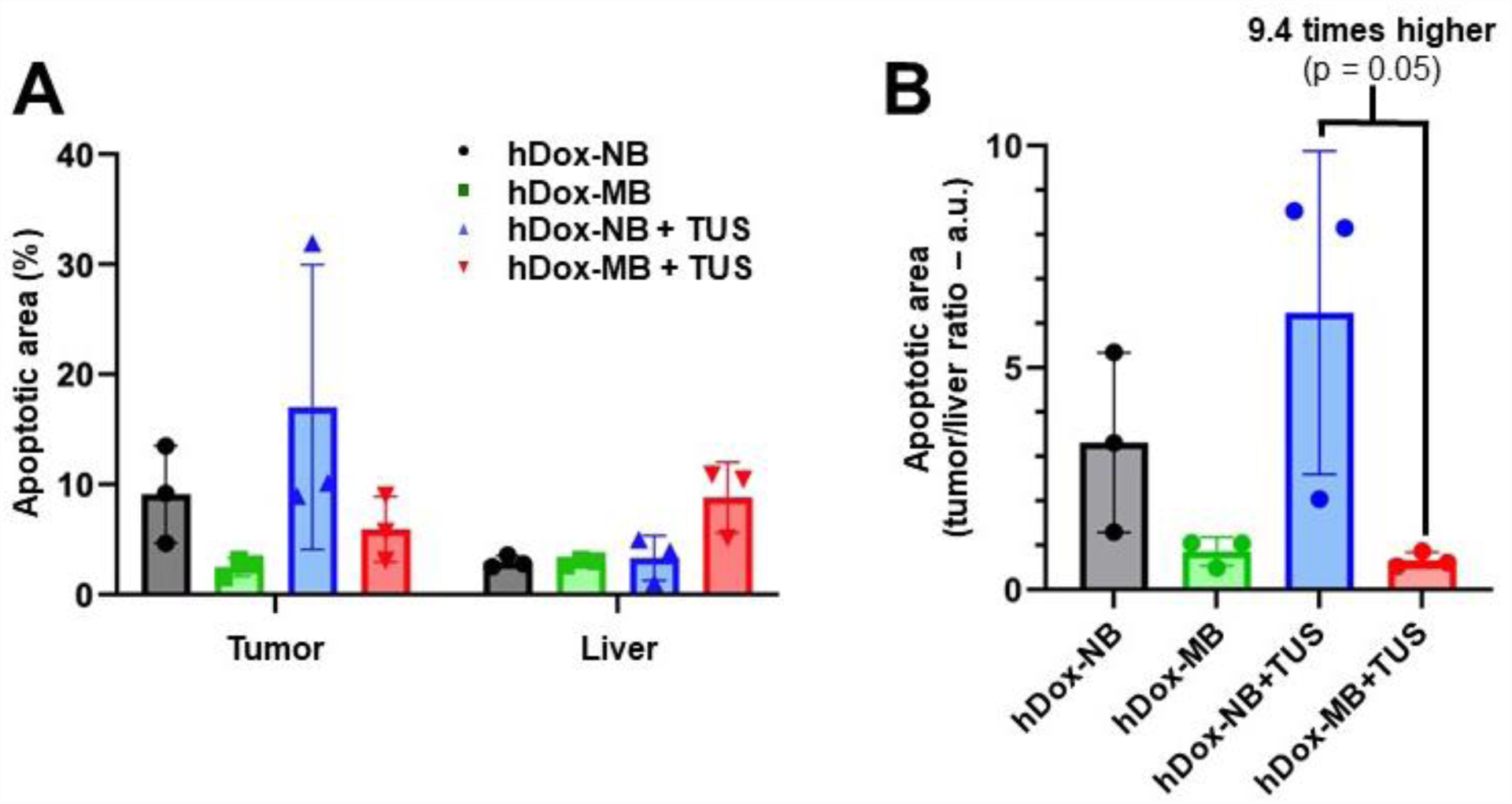
The apoptotic area calculated from TUNEL staining. Percentages of TUNEL-positive area in tumors and livers were quantified using the Positive Pixel Count algorithm in the Image-Scope analysis software (A); Tumor-to-liver ratio of the apoptotic area (B). Data are expressed as mean ± standard deviations (n = 3).

## Discussion

The use of acoustically active micro- and nanobubbles in conjunction with low frequency (1-3 MHz) and higher intensity (0.3-3 W/cm^2^) US in cancer therapy has been shown in many studies to improve drug delivery efficiency and elicit potential synergistic effects when combined with other therapeutic modalities, such as chemotherapy or immunotherapy (32,35,38,44,62). Despite numerous assertions that drug delivery to solid tumors could be more effective using NBs compared to MBs due to smaller bubble diameters, the effect of bubble size on drug delivery efficiency to tumors has not been previously examined directly. Accordingly, this study, for the first time, rigorously compared the efficiency of ultrasound-mediated drug delivery using doxorubicin-loaded lipid-shelled NBs and MBs produced using identical shell materials and matched drug doses. Free drug was removed in the formulation to allow focus on drug-loaded bubbles only, and care was taken to remove as many sub-micron bubbles from the MB population as possible. For clinical relevance, the study was carried out in orthotopic liver tumors inoculated in immunocompetent rats.

We successfully prepared hDox-NBs and size-isolated hDox-MBs and purified them to remove any free drugs. While the gas and shell composition of hDox-NBs and hDox-MBs is the same, they are distinct in size, concentration, DLC, and gas volume. RMM and DLS confirmed hDox-NBs as 280 ± 123 nm and hDox-MBs as 1104 ± 373 nm (Figure 2A-B). Due to purification and size isolation procedures, hDox-MB final concentration (3.18x10^9^ ± 2.9x10^8^ MBs/mL) was 12 times lower than hDox-NBs (4.06x10^10^ ± 6.9x10^8^ NBs/mL). This MB concentration is comparable to commercial MBs (Lumason®, Definity®, Optison™) in the range of 3-4x10^8^ MBs/mL (63). To retain all shell components during washing, a mixture of PBS/Glycerol/PG was used. The drug loading content (DLC) of hDox-NBs (151.97 ± 7.6 µg/mL) was higher than hDox-MBs (131.11 ± 15.3 µg/mL), as shown in Figure 2C. When comparing DLC per bubble, hDox-MBs are 12 times higher than hDox-NBs (Figure 2D). Additionally, hDox-MBs possess more gas volume and a 15.6 times larger surface area than hDox-NBs. This indicates that NBs offer greater drug loading efficiency per unit surface area. The gas volume is important for acoustic activity comparison. NBs and MBs are typically evaluated under normalized gas volume conditions to ensure a relevant comparison (64,64). However, in the *in vivo* experiments, we administered an equal Dox dose (327.75 µg/kg) to examine the efficacy of US-mediated drug-loaded NBs compared to MBs.

The perfusion dynamic of hDox-NBs and hDox-MBs in a rat tumor was carried out to determine the start time and duration of TUS irradiation for maximal NBs therapeutic efficiency. Understanding the kinetics of perfusion of different-sized drug-loaded bubbles in vivo can provide useful information for developing more effective treatment strategies that could enhance therapeutic efficacy and ultimately improve treatment outcomes. Here, the timing of TUS irradiation is crucial for properly using the sensitizing capability of NBs outside of the vasculature. Theoretically, since the advantage of NBs is their ability to extravasate and accumulate in the tumor microenvironment, the TUS application should ideally be carried out after the bubbles have extravasated in the tumor. However, in this study, for consistency with MBs, we applied the TUS 2 min after bubble injection. At this point, NBs are likely in the tumor’s small vessels or at the beginning of their extravasation, although determining the exact stage of extravasation may be difficult. Additional studies should be conducted to understand whether extravascular or intravascular bubble cavitation, or a combination of both, is most effective in eliciting a robust therapeutic response. Ultimately, ensuring the presence of bubbles at the tumor site prior to TUS can increase the possibility of enhanced drug delivery and therapeutic effects.

By analyzing the ultrasound imaging nonlinear TIC, we found that both hDox-NBs and hDox-MBs exhibit a rapid increase in signal enhancement and reach the peak enhancement within 1 min post-injection (Figure 3). This finding was similar to the previous work where the pharmacokinetics of non-drug-loaded NBs were compared to Definity MBs in subcutaneous human colorectal adenocarcinoma (LS174T) in mice (37). They also showed that the decay rate of NBs in the tumor was slower than with MBs. In contrast, here, hDox-MBs were slightly more stable in the tumor than hDox-NBs under our experimental conditions, as indicated by a longer half-life of 7 min of MBs and a slower washout rate vs 2 min of NBs. The primary difference, compared to prior studies, is the bubble composition. Our MBs are comprised of the same material as NBs. Accordingly, the stability of the agents differs. In addition, because we matched the injected drug dose between MBs and NBs, the difference in gas volume and bubble concentration influences the signal decay rate. Signal decay rates of these bubbles could also be related to clearance by the reticuloendothelial system (RES) or potential extravasation (65–71). Based on these findings, we initiated TUS irradiation 2 minutes after bubble administration. This timing was chosen to coincide with the washout phase, where both hDox-MBs and hDox-NBs concentration in the tumor is approximately 80% over the peak enhancement, driving enough time for NB to begin extravasate. We irradiated the tumor with TUS for 5 min, corresponding to 15% and 50% over the peak enhancement of the remaining NBs and MBs, respectively.

The biodistribution at 3 hours after treatment was evaluated semi-quantitatively by comparing the mean fluorescence intensity of Dox in tumors and organs obtained from optical fluorescence imaging. While no significant differences emerged in biodistribution, noticeable trends were apparent in the results. Considering the distribution within organs, the prominent finding is the absence of drug fluorescence in the heart and only minimal intensity in the lungs, regardless of the treatment groups. Both NBs and MBs displayed generally higher Dox signal within the tumor compared to other organs except for the spleen (Figure 4B). In general, a significant portion of nanoparticles tends to be sequestered within the liver (15 – 200 nm) and spleen (200 – 400 nm) or eliminated through the renal system (<15 nm) upon administration into the body (72). The size of the particles is recognized as one of the crucial factors that correlates with the rate of uptake and retention within cells (73).

Interestingly, our findings revealed an increased signal of Dox in the liver for hDox-MB+TUS, whereas the opposite trend was observed for NBs. Furthermore, the application of TUS appeared to minimize uptake within the spleen. This phenomenon could be attributed to the transformation of smaller particles from MBs due to the influence of cavitation. It could also be attributed to lipid-based UCAs being rapidly captured by macrophages and cleared via the reticular endothelial system (RES) (68,70,71,74). It has been reported that particle larger than 100 – 200 nm tends to be removed by splenic filtration within one to four hour after particle administration (72,75,76). Therefore, sequestered bubbles or lipid fragments with Dox could possibly be found in the spleen. Particularly noteworthy is the substantial Dox signal associated hDox-NB, indicating the effective delivery of Dox to the tumor area. Studies showed that nanoparticles within the 40 to 400 nm size range exhibit increased tumor accumulation while minimizing renal clearance (77,78). In contrast, hDox-MBs alone displayed lower signals in the tumors compared to the liver, highlighting the influence of size on the tumor targeting process. This implies that NBs exhibited higher tumor signal even without TUS treatment, highlighting their superior performance in tumor targeting.

Focusing on quantitative drug amounts within the tumor tissue (Figure 4C), a noticeable finding is the narrower variance among hDox-NBs samples, indicating a more consistent response facilitated by smaller bubbles. Moreover, a clear effect of TUS was observed with hDox-NBs+TUS, resulting in 3-fold increased cellular uptake. In contrast, this effect decreased with hDox-MBs, possibly because Dox access to the tumor area with MBs, whether with or without TUS, mainly relied on the liver’s enhanced permeability and retention (EPR) effect. This mechanism also accounts for the notable variability among MB samples.

Ultrasound-triggered MBs destruction through inertial cavitation has been reported to induce various vascular bioeffects (1–3), including high shear stress, elevated temperatures, micro-jetting, and invagination of micro-vessels (4). Consequently, these bioeffects can lead to microvascular damage, hemorrhage, endothelial cell injury, and apoptosis (5–7). In previous studies, NBs also have been explored as vascular-disrupting agents, showing similar impacts as MBs on tumor vascularity and potential damage to blood vessels through inertial cavitation near endothelial cells (8). Although MBs can cause increased vascular permeability, amplify tumoral drug release, induce thrombolysis of blood clots, and even open the blood-brain barrier under US stimulation, they primarily remain within the intravascular space (9,10). In contrast, NBs have demonstrated the ability to escape tumor vasculature because the vessels are poorly formed and have pathologically large endothelial gaps, enabling their use to directly target tumor cells and increase therapeutic agent uptake (11–14).

The immunohistochemistry (IHC) analysis of tumor sections using the TUNEL assay provided valuable insights into the therapeutic efficiency and acute damage in the tumor and surrounding liver tissue. The TUNEL assay detects DNA fragmentation, indicating apoptotic cell death (79,80). In this study, representative TUNEL slides of tumors in the liver were examined. Our TUNEL analysis of apoptotic cell death demonstrated distinct patterns among the treatment groups. The hDox-NBs without TUS (Figure 5A) exhibited a higher percentage of apoptosis in tumors compared to hDox-MBs (Figure 5B), aligning with our biodistribution findings. This can be attributed to the ability of smaller NBs to passively extravasate from tumor vessels into the tumor interstitium, allowing them to interact closely with tumor cells before cavitation and maximize drug delivery (36,81). Remarkably, even with similar total drug accumulation, hDox-NB+TUS treatment exhibited a significantly larger apoptotic cell death area within the tumor and fewer off-target apoptotic cells in the normal liver (Figure 5C) compared to hDox-MB+TUS treatment (Figure 5D). The percentage of apoptotic cells within the region of interest was notably higher in tumors treated with hDox-NB+TUS compared to hDox-MB+TUS, displaying a 2.8-fold increase (Figure 6A). This trend was further validated by the significantly elevated tumor-to-liver apoptotic ratio following hDox-NB+TUS treatments compared to hDox-MB+TUS treatments (Figure 6B).

The enhanced therapeutic efficacy of hDox-NB+TUS in inducing tumor-specific apoptosis could be attributed to the NBs’ potential sensitizing ability when cavitated outside the vasculature, intensifying cytotoxic effects and triggering apoptotic cell death within the tumor microenvironment (38). Notably, NBs preferentially accumulate in the tumor’s smaller vessels and extravascular space, minimizing off-target effects. In contrast, MBs predominantly reside within vessels, including those in normal tissue, limiting their potential to induce cell death in the distant tumor parenchyma away from these vessels. Due to their intravascular residence, MBs are more likely to cause vascular damage and collapse upon cavitation (82,83). Furthermore, the divergent therapeutic effects of NBs and MBs may also be linked to concentration-dependent phenomena. Our findings indicated that NBs induced apoptosis primarily at higher concentrations, localized within the tumor microenvironment where they predominantly accumulate. In contrast, MBs exhibited significant effects even at lower concentrations, potentially resulting in off-target impacts since they did not accumulate in the tumor. Overall, our results emphasize the enhanced specificity of hDox-NB+TUS in inducing apoptotic cell death within tumor tissue while minimizing damage to healthy liver tissue, in contrast to the broader apoptosis impact observed with MBs across the entire organ. These findings are in line with the drug delivery accumulation without TUS, emphasizing the amplified effect within the tumor region with NBs likely due to their penetration capability, compared to MBs which had a considerably reduced impact when unable to access the tumor region.

Prior research has primarily examined MBs and NBs in flank tumor models (84–86), where human cell lines are implanted in the subcutaneous region of animals, allowing easy tumor monitoring. However, these models can differ immunologically and phenotypically from orthotopic models, where tumors naturally form (87,88). Subcutaneous tumors exhibit irregular vasculature with thin-walled vessels, while orthotopic tumors have more abundant vasculature with thicker walls, influencing drug delivery dynamics due to varying perfusion and hypoxia levels (89,90). Despite the prevalence of flank models, some researchers have explored MBs and NBs in orthotopic tumor models, recognized as optimal for studying therapeutic strategies (91–95). Our study employed an HCC orthotopic tumor model in a larger animal, making available a large area of normal tissue surrounding the cancerous lesions. Orthotopic tumor models offer several key benefits, including enhanced clinical relevance, accurate assessment of drug response, realistic drug distribution, and translational relevance. Furthermore, the rat model used in our study involved immunocompetent rats, which enhances the translational applicability of our findings by accounting for the immune response in a more clinically relevant manner (96–98). This approach enables a more precise evaluation of drug delivery in the liver tissue microenvironment.

This study has limitations. Firstly, despite the distinct average sizes of our NBs and MBs, a degree of contamination of NBs within our MB population is evident (Figure 2A). This inadvertent inclusion of NBs could potentially influence the outcomes attributed to the MBs. Secondly, we assessed acute damage in the tumor and adjacent liver tissue within 3 hours after treatment, aiming to capture the immediate effects of drug-loaded NBs in the tumor microenvironment. However, this timeframe might not entirely reflect the treatment’s long-term impact on tumor-related aspects like development, survival, immune response, drug resistance, metastasis, or treatment-related side effects. Longer time points are needed for assessing long-term therapeutic efficacy. Next, the sample size was small, with only five rats per group. Despite careful design and rigorous analysis, this limited sample size might reduce statistical power and generalizability (99). Further studies with larger samples are necessary to validate our findings.

Additionally, tumor size variation among groups (Figure S6) could have affected drug delivery and therapeutic outcomes. Despite efforts to minimize this variation, tumor size impacts factors like blood flow, vascular permeability, and drug penetration, influencing drug delivery efficacy (97). Interpretation and comparison of results across different tumor sizes require caution. Further studies with uniform tumor sizes or strategies to account for variations are needed. Lastly, we didn’t directly examine acoustic cavitation activity, which is important for inducing vascular bioeffects. Treatment parameters were based on prior studies (100–102). Evaluating acoustic parameters like pressure, frequency, exposure time, and bubble concentration on cavitation dose for NBs and MBs would be beneficial.

## Conclusion

In summary, the current study rigorously evaluated the delivery efficiency of drug-loaded NBs compared to MBs in orthotopic liver tumors. Our findings elucidate the district behavior of bubbles during an initial 3-hour time point following injection and sonication at 80% of the peak enhancement. At these conditions, both hDox-NBs and hDox-MBs have comparable delivery efficiency and similar biodistribution and pharmacokinetic characteristics. However, when coupled with TUS, hDox-NBs were shown to be much more effective than hDox-MB at delivering tumor-specific, normal tissue-sparing, highly lethal drug doses to cancer cells, as indicated by the increased degree of tumor damage observed following treatment with hDox-NB+TUS. These findings suggest that hDox-NBs could be a superior approach for treating tumors of the liver and possibly liver metastasis and other aggressive cancers. Based on this data, NBs can deliver higher concentrations of the chemotherapeutic agent directly to the tumor site while minimizing systemic toxicity. Additional study is needed to evaluate the contribution of extravascular NB cavitation, long-term efficacy of hDox-NBs, as well as to optimize their formulation and delivery parameters to maximize their therapeutic potential. Likewise, mechanistic studies examining the role of the immune system in delivery efficiency and therapeutic outcomes in these models are needed to fully appreciate the observed differences.

## Acknowledgements

This work was supported by the National Institutes of Health (Award no. 5R01EB028144-04) and Visual Sciences Research Center Core Facilities funded by NIH P30 core grant (EY011373). Amin Sojahrood was supported by Natural Sciences and Engineering Research Council of Canada (NSERC). Felipe Berg was supported by Marcos Lottenberg & Marcus Wolosker International Fellowship for Physicians Scientist. We would like to express our gratitude to Selva Jeganathan, Emily Budziszewski, and Felipe Matsunaga for their assistance with tumor inoculation. Additionally, we would like to acknowledge Siemens Healthineers for providing the ultrasound device used in this study. Illustrations were created using BioRender.com.

## Notes

### Competing Interest Statement

The authors have declared no competing interest.

## References

1. Llovet JM, Kelley RK, Villanueva A, Singal AG, Pikarsky E, Roayaie S, et al. Hepatocellular carcinoma. Nat Rev Dis Primer 2021 71. 2021 Jan;7(1):1–28.

2. Griscom JT, Wolf PS. Liver Metastasis. StatPearls. 2022 Sep;

3. Ruers T, Bleichrodt RP. Treatment of liver metastases, an update on the possibilities and results. Eur J Cancer. 2002 May;38(7):1023–33.

4. Bakrania A, Zheng G, Bhat M. Nanomedicine in Hepatocellular Carcinoma: A New Frontier in Targeted Cancer Treatment. Pharmaceutics. 2022 Jan;14(1):41.

5. Yang S, Cai C, Wang H, Ma X, Shao A, Sheng J, et al. Drug delivery strategy in hepatocellular carcinoma therapy. Cell Commun Signal. 2022 Dec;20(1):1–14.

6. Osei E, Al-Asady A. A review of ultrasound-mediated microbubbles technology for cancer therapy: a vehicle for chemotherapeutic drug delivery. J Radiother Pract. 2020 Sep;19(3):291–8.

7. Sitta J, Howard CM. Applications of Ultrasound-Mediated Drug Delivery and Gene Therapy. Int J Mol Sci 2021 Vol 22 Page 11491. 2021 Oct;22(21):11491.

8. Metkar SP, Fernandes G, Navti PD, Nikam AN, Kudarha R, Dhas N, et al. Nanoparticle drug delivery systems in hepatocellular carcinoma: A focus on targeting strategies and therapeutic applications. OpenNano. 2023 Jul;12:100159.

9. Couture O, Foley J, Kassell NF, Larrat B, Aubry JF. Review of ultrasound mediated drug delivery for cancer treatment: updates from pre-clinical studies. Transl Cancer Res. 2014 Oct;3(5).

10. Sirsi SR, Borden MA. Microbubble Compositions, Properties and Biomedical Applications. Bubble Sci Eng Technol. 2009;1(1–2):3.

11. Martin KH, Dayton PA. Current Status and Prospects for Microbubbles in Ultrasound Theranostics. Wiley Interdiscip Rev Nanomed Nanobiotechnol. 2013 Jul;5(4):329–45.

12. Paefgen V, Doleschel D, Kiessling F. Evolution of contrast agents for ultrasound imaging and ultrasound-mediated drug delivery. Front Pharmacol. 2015 Sep 15;6:197.

13. Kooiman K, Vos HJ, Versluis M, de Jong N. Acoustic behavior of microbubbles and implications for drug delivery. Adv Drug Deliv Rev. 2014 Jun 15;72:28–48.

14. Omid C. Farokhzad Juliana M. Chan RL Pedro M Valencia, Liangfang Zhang. Polymeric Nanoparticles for Drug Delivery Juliana. J Cancer Sci Ther. 2012;4(1):163–75.

15. Sharma D, Giles A, Hashim A, Yip J, Ji Y, Do NNA, et al. Ultrasound microbubble potentiated enhancement of hyperthermia-effect in tumours. PloS One. 2019;14(12):e0226475.

16. Ingram N, McVeigh LE, Abou-Saleh RH, Maynard J, Peyman SA, McLaughlan JR, et al. Ultrasound-triggered therapeutic microbubbles enhance the efficacy of cytotoxic drugs by increasing circulation and tumor drug accumulation and limiting bioavailability and toxicity in normal tissues. Theranostics. 2020;10(24):10973–92.

17. Wang TY, Wilson KE, Machtaler S, Willmann JK. Ultrasound and Microbubble Guided Drug Delivery: Mechanistic Understanding and Clinical Implications. Curr Pharm Biotechnol. 2014 Feb;14(8):743.

18. Tsutsui JM, Xie F, Porter RT. The use of microbubbles to target drug delivery. Cardiovasc Ultrasound. 2004 Nov 16;2(1):23.

19. Deprez J, Lajoinie G, Engelen Y, De Smedt SC, Lentacker I. Opening doors with ultrasound and microbubbles: Beating biological barriers to promote drug delivery. Adv Drug Deliv Rev. 2021 May 1;172:9–36.

20. Liu HL, Fan CH, Ting CY, Yeh CK. Combining Microbubbles and Ultrasound for Drug Delivery to Brain Tumors: Current Progress and Overview. Theranostics. 2014 Feb 12;4(4):432–44.

21. Al-Jawadi S, Thakur SS. Ultrasound-responsive lipid microbubbles for drug delivery: A review of preparation techniques to optimise formulation size, stability and drug loading. Int J Pharm. 2020 Jul 30;585:119559.

22. Ramirez DG, Abenojar E, Hernandez C, Lorberbaum DS, Papazian LA, Passman S, et al. Contrast-enhanced ultrasound with sub-micron sized contrast agents detects insulitis in mouse models of type1 diabetes. Nat Commun. 2020 May 7;11:2238.

23. Maeda H. The enhanced permeability and retention (EPR) effect in tumor vasculature: the key role of tumor-selective macromolecular drug targeting. Adv Enzyme Regul. 2001 May;41(1):189–207.

24. Maeda H. Toward a full understanding of the EPR effect in primary and metastatic tumors as well as issues related to its heterogeneity. Adv Drug Deliv Rev. 2015 Aug;91:3–6.

25. Jain RK, Stylianopoulos T. Delivering nanomedicine to solid tumors. Nat Rev Clin Oncol 2010 711. 2010 Sep;7(11):653–64.

26. Batchelor DVB, Armistead FJ, Ingram N, Peyman SA, Mclaughlan JR, Coletta PL, et al. Nanobubbles for therapeutic delivery: Production, stability and current prospects. Curr Opin Colloid Interface Sci. 2021 Aug;54:101456.

27. Leon AD, Perera R, Hernandez C, Cooley M, Jung O, Jeganathan S, et al. Contrast enhanced ultrasound imaging by nature-inspired ultrastable echogenic nanobubbles. Nanoscale. 2019 Aug;11(33):15647–58.

28. Perera RH, Abenojar E, Nittayacharn P, Wang X, Ramamurthy G, Peiris P, et al. Intracellular vesicle entrapment of nanobubble ultrasound contrast agents targeted to PSMA promotes prolonged enhancement and stability in vivo and in vitro. Nanotheranostics. 2022;6(3):270.

29. Batchelor DVB, Abou-Saleh RH, Coletta PL, McLaughlan JR, Peyman SA, Evans SD. Nested Nanobubbles for Ultrasound-Triggered Drug Release. ACS Appl Mater Interfaces. 2020 Jul;12(26):29085–93.

30. Nittayacharn P, Yuan HX, Hernandez C, Bielecki P, Zhou H, Exner AA. Enhancing Tumor Drug Distribution With Ultrasound-Triggered Nanobubbles. J Pharm Sci. 2019 Sep 1;108(9):3091–8.

31. Nittayacharn P, Abenojar E, Leon AD, Wegierak D, Exner AA. Increasing Doxorubicin Loading in Lipid-Shelled Perfluoropropane Nanobubbles via a Simple Deprotonation Strategy. Front Pharmacol. 2020 May;11:644.

32. Exner AA, Kolios MC. Bursting microbubbles: How nanobubble contrast agents can enable the future of medical ultrasound molecular imaging and image-guided therapy. Curr Opin Colloid Interface Sci. 2021 Aug;54:101463.

33. Jin J, Yang L, Chen F, Gu N. Drug delivery system based on nanobubbles. Interdiscip Mater. 2022 Oct;1(4):471–94.

34. Cavalli R, Soster M, Argenziano M. Nanobubbles: a promising efficient tool for therapeutic delivery. Ther Deliv. 2016 Feb;7(2):117–38.

35. Bharti K, Kumar M, Jha A, Mishra B. Nanobubbles to aid drug delivery. Syst Nanovesicular Drug Deliv. 2022 Jan;323–36.

36. Wu H, Abenojar EC, Perera R, De Leon AC, An T, Exner AA. Time-intensity-curve Analysis and Tumor Extravasation of Nanobubble Ultrasound Contrast Agents. Ultrasound Med Biol. 2019 Sep 1;45(9):2502–14.

37. Wu H, Rognin NG, Krupka TM, Solorio L, Yoshiara H, Guenette G, et al. Acoustic Characterization and Pharmacokinetic Analyses of New Nanobubble Ultrasound Contrast Agents. Ultrasound Med Biol. 2013 Nov 1;39(11):2137–46.

38. Lu S, Zhao P, Deng Y, Liu Y. Mechanistic Insights and Therapeutic Delivery through Micro/Nanobubble-Assisted Ultrasound. Pharmaceutics. 2022 Mar;14(3):480.

39. Helbert A, Gaud E, Segers T, Botteron C, Frinking P, Jeannot V. Monodisperse versus Polydisperse Ultrasound Contrast Agents: In Vivo Sensitivity and safety in Rat and Pig. Ultrasound Med Biol. 2020 Dec;46(12):3339–52.

40. Shekhar H, Smith NJ, Raymond JL, Holland CK. Effect of Temperature on the Size Distribution, Shell Properties, and Stability of Definity®. Ultrasound Med Biol. 2018 Feb 1;44(2):434–46.

41. Faez T, Goertz D, De Jong N. Characterization of Definity^TM^ Ultrasound Contrast Agent at Frequency Range of 5–15 MHz. Ultrasound Med Biol. 2011 Feb 1;37(2):338–42.

42. Zhu X, Guo J, He C, Geng H, Yu G, Li J, et al. Ultrasound triggered image-guided drug delivery to inhibit vascular reconstruction via paclitaxel-loaded microbubbles. Sci Rep 2016 61. 2016 Feb;6(1):1– 12.

43. Shuai X, Ai H, Nasongkla N, Kim S, Gao J. Micellar carriers based on block copolymers of poly(ε-caprolactone) and poly(ethylene glycol) for doxorubicin delivery. J Controlled Release. 2004 Aug;98(3):415–26.

44. Wang M, Zhang Y, Cai C, Tu J, Guo X, Zhang D. Sonoporation-induced cell membrane permeabilization and cytoskeleton disassembly at varied acoustic and microbubble-cell parameters. Sci Rep 2018 81. 2018 Mar;8(1):1–12.

45. Lentacker I, Geers B, Demeester J, Smedt SCD, Sanders NN. Design and Evaluation of Doxorubicin-containing Microbubbles for Ultrasound-triggered Doxorubicin Delivery: Cytotoxicity and Mechanisms Involved. Mol Ther. 2010 Jan;18(1):101–8.

46. Feshitan JA, Chen CC, Kwan JJ, Borden MA. Microbubble size isolation by differential centrifugation. J Colloid Interface Sci. 2009 Jan;329(2):316–24.

47. Abenojar EC, Nittayacharn P, Leon ACD, Perera R, Wang Y, Bederman I, et al. Effect of Bubble Concentration on the in Vitro and in Vivo Performance of Highly Stable Lipid Shell-Stabilized Micro- And Nanoscale Ultrasound Contrast Agents. Langmuir. 2019 Aug;35(31):10192–202.

48. Choi B, Pe J, Yu B, Kim DH. Syngeneic N1-S1 Orthotopic Hepatocellular Carcinoma in Sprague Dawley Rat for the Development of Interventional Oncology-Based Immunotherapy: Survival Assay and Tumor Immune Microenvironment. Cancers. 2023 Feb;15(3):913.

49. Chan HH, Chu TH, Chien HF, Sun CK, Wang EM, Pan HB, et al. Rapid induction of orthotopic hepatocellular carcinoma in immune-competent rats by non-invasive ultrasound-guided cells implantation. BMC Gastroenterol. 2010 Jul;10(1):1–11.

50. Lee TK, Na KS, Kim J, Jeong HJ. Establishment of Animal Models with Orthotopic Hepatocellular Carcinoma. Nucl Med Mol Imaging. 2014;48(3):173.

51. Riccabona M, Nelson TR, Pretorius DH, Davidson TE. Distance and volume measurement using three-dimensional ultrasonography. J Ultrasound Med Off J Am Inst Ultrasound Med. 1995;14(12):881–6.

52. Bleuzen A, Tranquart F. Incidental liver lesions: diagnostic value of cadence contrast pulse sequencing (CPS) and SonoVue. Eur Radiol. 2004 Oct;14 Suppl 8:P53–62.

53. de Leon A, Wei P, Bordera F, Wegierak D, McMillen M, Yan D, et al. Pickering Bubbles as Dual-Modality Ultrasound and Photoacoustic Contrast Agents. ACS Appl Mater Interfaces. 2020 May 13;12(19):22308–17.

54. Phillips PJ. Contrast pulse sequences (CPS): imaging nonlinear microbubbles. In: 2001 IEEE Ultrasonics Symposium Proceedings An International Symposium (Cat No01CH37263). 2001. p. 1739– 45 vol.2.

55. Forsberg F, Shi WT, Merritt CRB, Dai Q, Solcova M, Goldberg BB. On the Usefulness of the Mechanical Index Displayed on Clinical Ultrasound Scanners for Predicting Contrast Microbubble Destruction. J Ultrasound Med. 2005;24(4):443–50.

56. Jeganathan S, Budziszewski E, Bielecki P, Kolios MC, Exner AA. In situ forming implants exposed to ultrasound enhance therapeutic efficacy in subcutaneous murine tumors. J Controlled Release. 2020 Aug;324:146–55.

57. Jeganathan S, Budziszewski E, Hernandez C, Dhingra A, Exner AA. Improving Treatment Efficacy of In Situ Forming Implants via Concurrent Delivery of Chemotherapeutic and Chemosensitizer. Sci Rep 2020 101. 2020 Apr;10(1):1–10.

58. Janke LJ, Ward JM, Vogel P. Classification, Scoring, and Quantification of Cell Death in Tissue Sections. Vet Pathol. 2019 Jan;56(1):33–8.

59. Dunn WD, Gearing M, Park Y, Zhang L, Hanfelt J, Glass JD, et al. Applicability of digital analysis and imaging technology in neuropathology assessment. Neuropathol Off J Jpn Soc Neuropathol. 2016 Jun;36(3):270.

60. Hernandez C, C. Abenojar E, Hadley J, Leon AC de, Coyne R, Perera R, et al. Sink or float? Characterization of shell-stabilized bulk nanobubbles using a resonant mass measurement technique. Nanoscale. 2019;11(3):851–5.

61. Milligan JJ, Saha S. A Nanoparticle’s Journey to the Tumor: Strategies to Overcome First-Pass Metabolism and Their Limitations. Cancers. 2022 Mar 29;14(7):1741.

62. Liu Y, Xie Q, Ma Y, Lin C, Li J, Hu B, et al. Nanobubbles containing PD-L1 Ab and miR-424 mediated PD-L1 blockade, and its expression inhibition to enable and potentiate hepatocellular carcinoma immunotherapy in mice. Int J Pharm. 2022 Dec 15;629:122352.

63. Characteristics and Echogenicity of Clinical Ultrasound Contrast Agents: An In Vitro and In Vivo Comparison Study - Hyvelin - 2017 - Journal of Ultrasound in Medicine - Wiley Online Library.

64. Abenojar EC, Bederman I, Leon AC de, Zhu J, Hadley J, Kolios MC, et al. Theoretical and Experimental Gas Volume Quantification of Micro- and Nanobubble Ultrasound Contrast Agents. Pharm 2020 Vol 12 Page 208. 2020 Mar;12(3):208.

65. Sirsi S, Feshitan J, Kwan J, Homma S, Borden M. Effect of Microbubble Size on Fundamental Mode High Frequency Ultrasound Imaging in Mice. Ultrasound Med Biol. 2010 Jun;36(6):935–48.

66. Sarkar K, Katiyar A, Jain P. Growth and dissolution of an encapsulated contrast microbubble: effects of encapsulation permeability. Ultrasound Med Biol. 2009 Aug;35(8):1385–96.

67. Borden MA, Longo ML. Dissolution Behavior of Lipid Monolayer-Coated, Air-Filled Microbubbles: Effect of Lipid Hydrophobic Chain Length. Langmuir. 2002 Nov 1;18(24):9225–33.

68. Morel DR, Schwieger I, Hohn L, Terrettaz J, Llull JB, Cornioley YA, et al. Human pharmacokinetics and safety evaluation of SonoVue, a new contrast agent for ultrasound imaging. Invest Radiol. 2000 Jan;35(1):80–5.

69. Dasgupta A, Sun T, Palomba R, Rama E, Zhang Y, Power C, et al. Nonspherical ultrasound microbubbles. Proc Natl Acad Sci. 2023 Mar 28;120(13):e2218847120.

70. Levchenko TS, Rammohan R, Lukyanov AN, Whiteman KR, Torchilin VP. Liposome clearance in mice: the effect of a separate and combined presence of surface charge and polymer coating. Int J Pharm. 2002 Jun 20;240(1):95–102.

71. Liu D, Hu Q, Song YK. Liposome clearance from blood: different animal species have different mechanisms. Biochim Biophys Acta BBA - Biomembr. 1995 Dec 13;1240(2):277–84.

72. Cataldi M, Vigliotti C, Mosca T, Cammarota M, Capone D. Emerging Role of the Spleen in the Pharmacokinetics of Monoclonal Antibodies, Nanoparticles and Exosomes. Int J Mol Sci. 2017 Jun 10;18(6):1249.

73. Zhang YN, Poon W, Tavares AJ, McGilvray ID, Chan WCW. Nanoparticle–liver interactions: Cellular uptake and hepatobiliary elimination. J Controlled Release. 2016 Oct 28;240:332–48.

74. Willmann JK, Cheng Z, Davis C, Lutz AM, Schipper ML, Nielsen CH, et al. Targeted microbubbles for imaging tumor angiogenesis: assessment of whole-body biodistribution with dynamic micro-PET in mice. Radiology. 2008 Oct;249(1):212–9.

75. Moghimi SM, Hedeman H, Muir IS, Illum L, Davis SS. An investigation of the filtration capacity and the fate of large filtered sterically-stabilized microspheres in rat spleen. Biochim Biophys Acta BBA - Gen Subj. 1993 Jun 11;1157(2):233–40.

76. Factors Controlling Nanoparticle Pharmacokinetics: An Integrated Analysis and Perspective | Annual Review of Pharmacology and Toxicology.

77. Liechty WB, Peppas NA. Expert Opinion: Responsive Polymer Nanoparticles in Cancer Therapy. Eur J Pharm Biopharm. 2012 Feb;80(2):241–6.

78. Subhan MA, Yalamarty SSK, Filipczak N, Parveen F, Torchilin VP. Recent Advances in Tumor Targeting via EPR Effect for Cancer Treatment. J Pers Med. 2021 Jun;11(6):571.

79. Loo DT. TUNEL assay. An overview of techniques. Methods Mol Biol Clifton NJ. 2002;203:21–30.

80. Yuan YG, Peng QL, Gurunathan S. Silver nanoparticles enhance the apoptotic potential of gemcitabine in human ovarian cancer cells: combination therapy for effective cancer treatment. Int J Nanomedicine. 2017 Sep 5;12:6487–502.

81. Karshafian R, Bevan PD, Williams R, Samac S, Burns PN. Sonoporation by Ultrasound-Activated Microbubble Contrast Agents: Effect of Acoustic Exposure Parameters on Cell Membrane Permeability and Cell Viability. Ultrasound Med Biol. 2009 May;35(5):847–60.

82. Wang J, Zhao Z, Shen S, Zhang C, Guo S, Lu Y, et al. Selective depletion of tumor neovasculature by microbubble destruction with appropriate ultrasound pressure. Int J Cancer. 2015 Nov 15;137(10):2478–91.

83. Padilla F, Brenner J, Prada F, Klibanov AL. Theranostics in the vasculature: bioeffects of ultrasound and microbubbles to induce vascular shutdown. Theranostics. 2023 Jul 14;13(12):4079– 101.

84. Al-Mahrouki AA, Iradji S, Tran WT, Czarnota GJ. Cellular characterization of ultrasound-stimulated microbubble radiation enhancement in a prostate cancer xenograft model. Dis Model Mech. 2014 Mar;7(3):363–72.

85. Marano F, Frairia R, Rinella L, Argenziano M, Bussolati B, Grange C, et al. Combining doxorubicin- nanobubbles and shockwaves for anaplastic thyroid cancer treatment: preclinical study in a xenograft mouse model. Endocr Relat Cancer. 2017 Jun;24(6):275–86.

86. Chiang PH, Fan CH, Jin Q, Yeh CK. Enhancing Doxorubicin Delivery in Solid Tumor by Superhydrophobic Amorphous Calcium Carbonate–Doxorubicin Silica Nanoparticles with Focused Ultrasound. Mol Pharm. 2022 Nov 7;19(11):3894–905.

87. Bibby MC. Orthotopic models of cancer for preclinical drug evaluation: advantages and disadvantages. Eur J Cancer. 2004 Apr 1;40(6):852–7.

88. Killion JJ, Radinsky R, Fidler IJ. Orthotopic Models are Necessary to Predict Therapy of Transplantable Tumors in Mice. Cancer Metastasis Rev. 1998 Sep 1;17(3):279–84.

89. Liu W, Zhu Y, Ye L, Zhu Y, Wang Y. Comparison of tumor angiogenesis in subcutaneous and orthotopic LNCaP mouse models using contrast-enhanced ultrasound imaging. Transl Cancer Res. 2021 Jul;10(7):3268–77.

90. Zhang W, Fan W, Rachagani S, Zhou Z, Lele SM, Batra SK, et al. Comparative Study of Subcutaneous and Orthotopic Mouse Models of Prostate Cancer: Vascular Perfusion, Vasculature Density, Hypoxic Burden and BB2r-Targeting Efficacy. Sci Rep. 2019 Jul 31;9(1):11117.

91. Kotopoulis S, Delalande A, Popa M, Mamaeva V, Dimcevski G, Gilja OH, et al. Sonoporation- enhanced chemotherapy significantly reduces primary tumour burden in an orthotopic pancreatic cancer xenograft. Mol Imaging Biol. 2014 Feb;16(1):53–62.

92. Optimal treatment occasion for ultrasound stimulated microbubbles in promoting gemcitabine delivery to VX2 tumors - PubMed.

93. Huang KW, Huang YC, Tai KF, Chen BH, Lee PH, Hwang LH. Dual Therapeutic Effects of Interferon-α Gene Therapy in a Rat Hepatocellular Carcinoma Model With Liver Cirrhosis. Mol Ther J Am Soc Gene Ther. 2008 Oct;16(10):1681–7.

94. Haram M, Snipstad S, Berg S, Mjønes P, Rønne E, Lage J, et al. Ultrasound and Microbubbles Increase the Uptake of Platinum in Murine Orthotopic Pancreatic Tumors. Ultrasound Med Biol. 2023 May;49(5):1275–87.

95. Evaluation of Loading Strategies to Improve Tumor Uptake of Gemcitabine in a Murine Orthotopic Bladder Cancer Model Using Ultrasound and Microbubbles - PubMed.

96. Ii REN, Jr JHR, Robertson JL, Arena CB, Davis EM, Singh RN, et al. Improved Local and Systemic Anti-Tumor Efficacy for Irreversible Electroporation in Immunocompetent versus Immunodeficient Mice. PLOS ONE. 2013 May 24;8(5):e64559.

97. Saini M, Bellinzona M, Meyer F, Cali‘ G, Samii M. Morphometrical Characterization of Two Glioma Models in the Brain of Immunocompetent and Immunodeficient Rats. J Neurooncol. 1999 Mar 1;42(1):59–67.

98. Buqué A, Galluzzi L. Modeling Tumor Immunology and Immunotherapy in Mice. Trends Cancer. 2018 Sep 1;4(9):599–601.

99. Andrade C. Sample Size and its Importance in Research. Indian J Psychol Med. 2020 Jan 6;42(1):102–3.

100. Wood AKW, Schultz SM, Lee WMF, Bunte RM, Sehgal CM. Antivascular ultrasound therapy extends survival of mice with implanted melanomas. Ultrasound Med Biol. 2010 May;36(5):853–7.

101. Wood A, Bunte R, Price H, Deitz M, Tsai J, Lee W, et al. The Disruption of Murine Tumor Neovasculature by Low-intensity Ultrasound – comparison between 1 MHz and 3 MHz sonication frequencies. Acad Radiol. 2008 Sep;15(9):1133–41.

102. Hunt SJ, Gade T, Soulen MC, Pickup S, Sehgal CM. Antivascular ultrasound therapy: magnetic resonance imaging validation and activation of the immune response in murine melanoma. J Ultrasound Med Off J Am Inst Ultrasound Med. 2015 Feb;34(2):275–87.

