## Supplementary Information for "Efficient ultrasound-mediated drug delivery to orthotopic liver tumors – Direct comparison of doxorubicin-loaded nanobubbles and microbubbles"

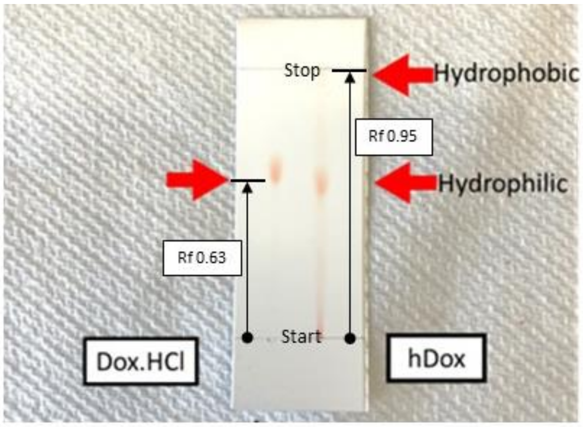


**Figure S1.** TLC plate showing the state of hydrophilic (left lane) and hydrophobic Dox (right lane). The Dox.HCl or hydrophilic Dox presented one spot on the TLC plate with an Rf of 0.63 while two spots with a larger Rf of 0.95 was found for the hDox. The ability of hDox to travel further on the TLC plate confirmed that it has less polarity (more hydrophobic).

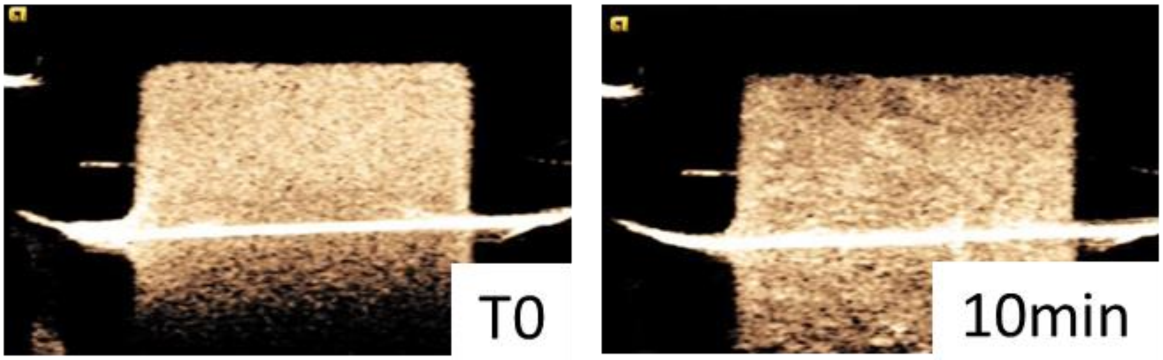


**Figure S2.** Representative ultrasound contrast images of purified hDox-NB at a 1:10 dilution with PBS (4.06 x$10^9$ NB/mL). The NB enhancement in the tissue-mimicking agarose phantom was imaged at 8 MHz with an 18 MHz probe, 0.1 MI. The left and right images showed the initial contrast and signal after 10 min of continuous scanning.

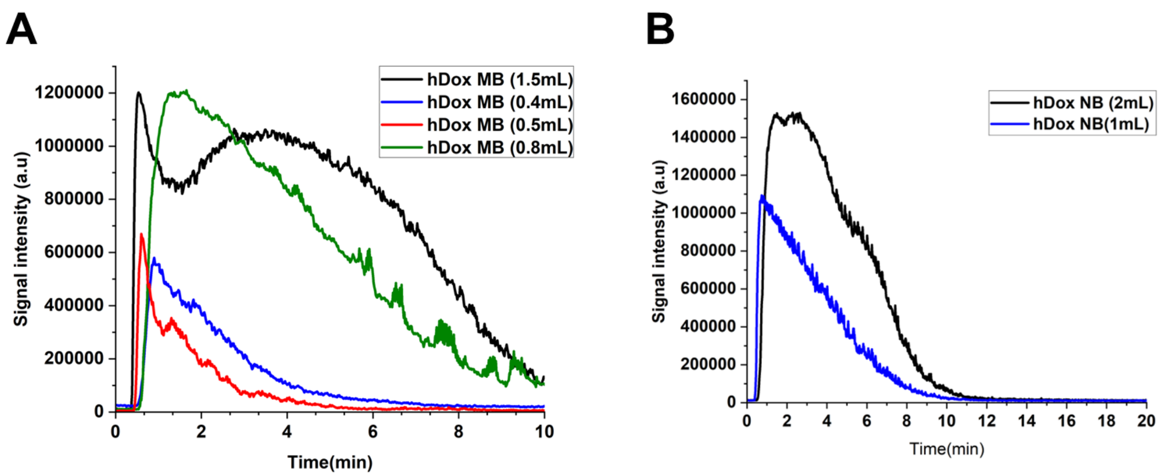


**Figure S3.** Time-intensity curves of tumor after bubble administration. hDox-MB for different injection volumes injected into the rats. Potential signal attenuation was present when 1.5 mL of hDox-MB was injected into the rat, as demonstrated by the rapid initial decrease in signal intensity over the first 2 minutes. Accordingly, to avoid tissue attenuation, the injection volume in this study was chosen to be below 1.5 mL (A). hDox-NB at 1 and 2 mL of injection volumes were injected into the rats (B).

**Table S1.** Summary of quantitative kinetic parameters of tumor obtained from time-intensity curve (TIC)

| Sample | Time to peak (min) | Peak Enhancement (a.u.) | Total AUC | Wash-in AUC | Washout AUC |
|---|---|---|---|---|---|
| hDox-MBs | 0.8 | 1251051.0 | 7557550.9 | 409248.3 | 7148302.6 |
| hDox-NBs | 0.8 | 1094257.0 | 4207794.5 | 309655.3 | 3898139.2 |
| % Difference between MB/NB | 0 | 14.33 | 79.61 | 32.16 | 93.38 |

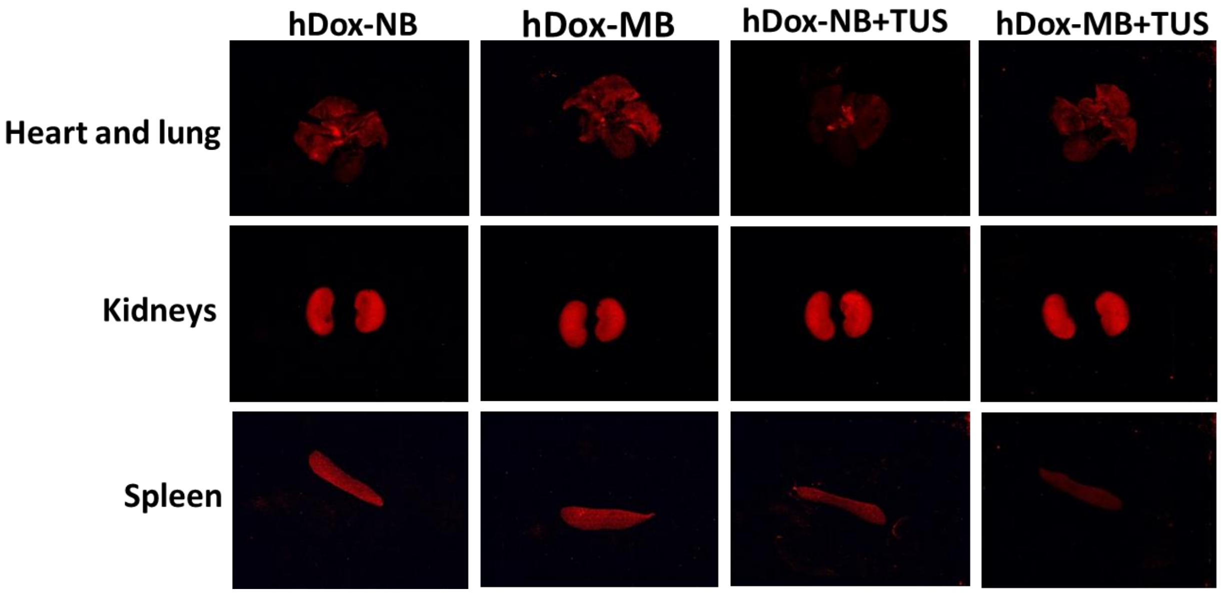


**Figure S4.** Representative optical fluorescence images of rat organs from each treatment group.

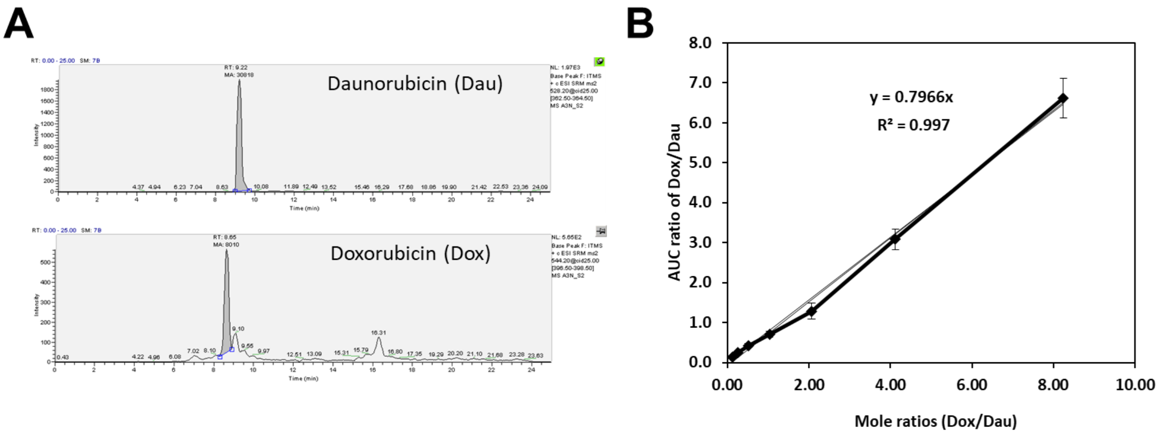


**Figure S5.** LC/MS spectra of a sample of rat's tumor showing chromatographic separation and LC/MS analysis of internal standard (Daunorubicin, Dau) and analyte (Doxorubicin, Dox) (A). Standard curve of Doxorubicin was prepared in tumor tissue homogenates by adding different amounts of Doxorubicin and Daunorubicin to reach final concentrations ranging from 5.4 to 43.5 ng/mL(B).

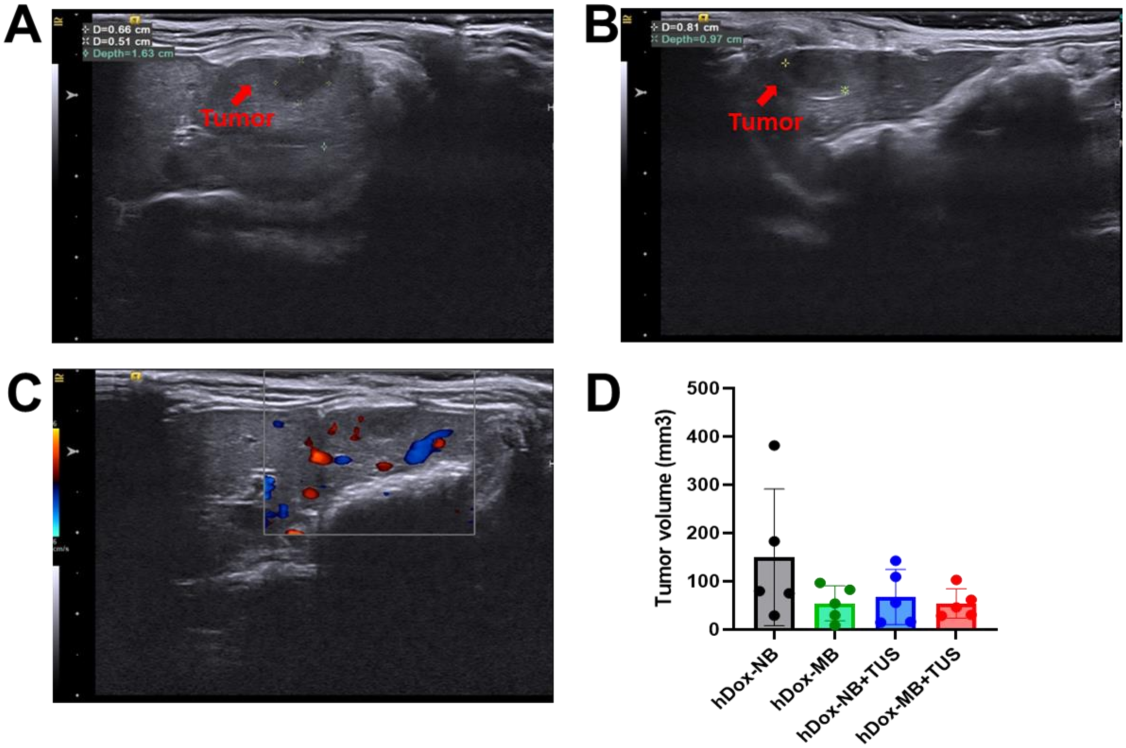


**Figure S6.** Representative B-mode images with tumor (red arrow) axis measurements. Volume was estimated by the standard clinical ellipsoidal equation, where V = 4/3π *a/2*b/2*c/2. Images illustrate the axial plane (A), sagittal plane (B), sagittal plane with color Doppler showing vascularized tumor (C), and average tumor volume in each treatment group (D). Data are presented as mean ± SD (n = 5).
